# Notch coordinates self-organization of germ layers and axial polarity in cnidarian gastruloids

**DOI:** 10.1101/2025.09.09.675057

**Authors:** Sanjay Narayanaswamy, Franziska Haas, Emmanuel Haillot, Elly Tanaka, Ulrich Technau

## Abstract

Dissociation and reaggregation experiments in several animal systems have revealed the stunning capacity of self-organization. Reaggregated early gastrula cells (here called gastruloids) of the sea anemone *Nematostella vectensis* are able to regenerate with only little delay whole organisms that are virtually indistinguishable from normal developing polyps. However, the molecular control underlying the restoration of body axis and germ layers remains largely unknown. To address this, we established a standardized protocol, which reproducibly generates gastruloids developing into polys with a single body axis. Here, we show that committed mesodermal and endodermal cells are sorted out to the surface of the aggregate, where mesodermal cells form clusters of about 30 cells. At a critical time point, one mesodermal cluster immigrates, along with peripherally attached endodermal cells. Thereby, the inner germ layer and the oral-aboral axis is established in one and the same process. Functional studies demonstrated that this polarization of the endodermal cells requires a feedback loop of Notch and Wnt signaling. The dissociation of the early embryo disrupts Notch signaling in the endodermal cells, which leads to transient adoption of an endomesodermal profile, marked by the expression of the mesodermal cadherin1 until the boundaries between the germ layer identities are re-established. Our results highlight distinct morphogenetic behaviors of mesodermal and endodermal cells and the hitherto unknown role of Notch signaling in germ layer boundary formation in self-organizing gastruloids. Conservation of Notch-mediated boundary formation between endoderm and mesoderm mirrors bilaterian mechanisms, demonstrating how adoption of ancestral regulatory networks governing morphogenesis likely enabled the diversification of metazoan body plans.

## Introduction

The establishment of body axes governs morphological diversity across 600 million years of animal radiation. A key step preceding body axes formation is tissue symmetry breaking. In many model organisms such as *C. elegans*, frogs and fruit flies, this event already occurs prior to or during fertilization and therefore, there are limited tractable systems to study spontaneous symmetry breaking in a tissue (Ishihara and Tanaka, 2018). Additionally, while conserved molecular feedback loops like between Wnt/β-catenin signaling and T-box transcription factors underpin axial patterning (Schwaiger et al. 2022; Anlas and Trivedi, 2021), their interplay with species-specific embryonic geometries and extraembryonic cues remains poorly understood. This knowledge gap stems from inherent limitations in studying native embryos, where maternal determinants and biomechanical constraints obscure cell-autonomous patterning principles (Ishihara and Tanaka, 2018). *Ex vivo* and *in vitro* self-organizing systems, particularly gastruloids, have emerged as excellent systems to dissect these mechanisms. By recapitulating axis formation through self-organization rather than preprogrammed embryonic architecture, these models enable us to isolate intrinsic modes of self-organization. For instance, human and mouse gastruloids can autonomously pattern their body axes mirroring the embryo, but without any inputs from the visceral endoderm or trophoblast. This highlights cell-autonomous symmetry breaking dependent on Wnt/Nodal signaling interactions rather than by external BMP gradients (Turner et al. 2017). Additionally, unlike in embryos, gastruloids pattern in minimal biochemical and mechanical constraints, allowing us to uncover conserved developmental models across species by removing species specific signaling and geometric constraints (Anlas and Trivedi, 2021). This explains why despite having radically different embryonic geometries, in vitro systems often converge on similar axial elongation dynamics, suggesting an adaptation of conserved circuits rather than de novo innovations (Manuel, 2008). Finally, these systems allow for fine tuning various experimental parameters such as cell proportions, helping us study aspects like dependence of symmetry breaking time on proportion of T+ cell populations (Oriola et al. 2024), something harder to test *in vivo*.

Both in native embryos and gastruloids, symmetry breaking and axis elongation are driven by the presence of localized signaling centers or organizers. The concept of the organizer, first established by Spemann and Mangold in amphibians (Spemann and Mangold, 1924), has since emerged as a unifying principle in developmental biology, with functionally similar regions identified across metazoans. This includes cnidarians, where earlier seminal transplantation experiments have led to the identification of the *Hydra* hypostome organizer (Browne, 1909; Broun and Bode, 2002; for reviews see Narayanaswamy and Technau, 2025; Galliot and Wenger, 2025), and more recently to the blastopore lip organizer in *Nematostella* (Kraus et al. 2016). Comparative studies revealed that the molecular logic underlying embryonic organizer function in *Nematostella* is deeply conserved across Metazoa. The Wnt/β-catenin signaling pathway, central to organizer activity in *Nematostella* as well as other cnidarians (Technau and Bode, 1999; Technau et al. 2000; Hobmayer et al., 2000; Momose et al., 2008; Duffy et al., 2010; Hayward et al., 2015; Kraus et al. 2016), also governs primary axis specification in bilaterians, where organizers such as the amphibian Spemann organizer and the mammalian node, direct embryonic patterning through localized Wnt signaling (Martinez Arias and Steventon, 2018). In *Nematostella*, transplantation experiments and targeted activation of Wnt signaling demonstrate that co-expression of *Wnt1* and *Wnt3* can confer organizer properties to ectopic regions, paralleling the axis-inducing capacity of organizer tissues in bilaterians. This highlights the deep evolutionary conservation of Wnt/β-catenin signaling in axial patterning, predating the split between cnidarians and bilaterians (Kraus et al. 2016; Lebedeva et al. 2021).

Cnidarians also possess remarkable regenerative capacities, with many organisms in this phylum possessing the ability to reestablish their body plan from dissociated and reaggregated cells from embryonic stages through to adult polyps, thus exhibiting a form of extreme regeneration (Gierer et al., 1972; Technau and Holstein, 1992; Technau et al. 2000; Kirillova and Kraus, 2018; for review see Narayanaswamy and Technau, 2025). While this phenomenon has been observed in various cnidarian polyps, the lack of accessible embryonic stages in most cnidarians has limited comparisons to early embryonic self-organization in bilaterians. In this regard, the anthozoan *Nematostella vectensis* stands out as a tractable model: its embryonic cells, when dissociated and reaggregated, can reestablish its primary body axis and germ layer topology, ultimately giving rise to a viable polyp indistinguishable from those developed through unperturbed embryogenesis. This plasticity is orchestrated by the *Wnt1* and *Wnt3* expressing organizer cells at the blastopore lip, whose presence is both necessary and sufficient for axis reestablishment in aggregates, mirroring the role of these organizers in normal development (Kraus et al. 2016; Kirillova et al. 2018). In addition, cells of the gastrodermis of the gastrulae become mesenchymal post-dissociation, followed by ingression in a manner similar to gastrodermal cells during embryonic development in hydrozoans. While the gastrodermal cells remember their fate in the aggregates, aboral ectodermal cells exhibit plasticity and can be reprogrammed into gastrodermal cells (Kirillova et al., 2018). Intriguingly, while *Nematostella* is morphologically diploblastic, recent work has shown that its germ layers exhibit molecular signatures homologous to the three germ layers of bilaterians where, the blastopore lip and resulting pharynx and septal filaments have an endodermal identity, while the gastrodermis correspond to the mesoderm (Steinmetz et al. 2017; Haillot et al. 2025). In this paradigm, the organizer cells reside within the endoderm tissue identity, reminiscent of the endodermal location of the Nieuwkoop center in amphibians (Gimlich, 1985; Gimlich and Gerhart, 1984). Yet, how this population of initially randomly distributed organizer cells clusters and undergoes a symmetry break in order to polarize the aggregate is poorly understood. This raises the quintessential question about the mechanism that underlies the self-organization of the organizer. Here we reveal the morphogenetic behavior and molecular identity of these germ layers in standardized gastruloids. We show that Notch signaling plays a crucial role in the polarization and symmetry break of the aggregate, which leads to the formation of the oral-aboral body axis as well as the topological segregation of the three germ layer identities in one step.

## Results

Previous methods used to make cell aggregates in *Nematostella* employed dissociation of gastrulae, followed by centrifugation and cutting the pellet by eye into pieces of roughly uniform size (Kirillova et al., 2018). There are some caveats with this method including the high degree of compaction leading to potentially large fraction of highly stressed/damaged cells, non-uniform sizes of aggregates leading to both variable number of axes as well as spatiotemporal expression patterns of genes. This variability makes it difficult to identify crucial molecular and cellular mechanisms underlying this process of self organization. To overcome this, we established a high throughput workflow (see Methods) **(Fig. 1A),** which generates standardized *Nematostella* gastruloids minimizing cell stress and giving rise to a specific axis number with reproducibility. We found that a seed count of 10,000 cells robustly gives rise to gastruloids that develop into polyps with a single axis **(Fig. 1B)**. Of note, we estimate that about 70% of seeded cells get incorporated into generating the gastruloid, based on the number of cells comprising the midgastrulae stage of the embryo (Kirillova et al. 2018). The remaining cells are expelled through a hitherto unknown mechanism. To examine the effect of seed count on the number of axes, aggregates of different seed counts were made using a *FoxA::mOrange2* reporter line, with the expression of fluorophore marking the future pharynx of the polyp. The number of fluorescent poles were then quantified across biological replicates as a proxy for the number of discrete axes formed. The results confirm a highly reproducible and scalable system of gastruloid generation **(Fig. 1C)**. Interestingly, based on the estimates of the number of blastopore lip cells present per embryo (Kirillova et al. 2018), this suggests a requirement of ∼155 blastopore lip cells (*Wnt1*/*Wnt3*+) per gastruloid to maintain the high axis inducing reproducibility of this system. To get further insights about axis scaling, we also examined tissue patterning by staining for markers endodermal and mesodermal markers *FoxA* and *SnailA* respectively expressed along the Oral-Aboral (OA) axis. We found that the gastruloids formed repeating modules of *FoxA* and *SnailA*+ domains proportional to the seed count, a pattern expected under a Turing-type reaction-diffusion model, where addition of more cells resolves into multiple repeating patterning modules dependent on cell number **(Fig. 1D)** (Ishihara and Tanaka, 2018).

**Figure 1.**
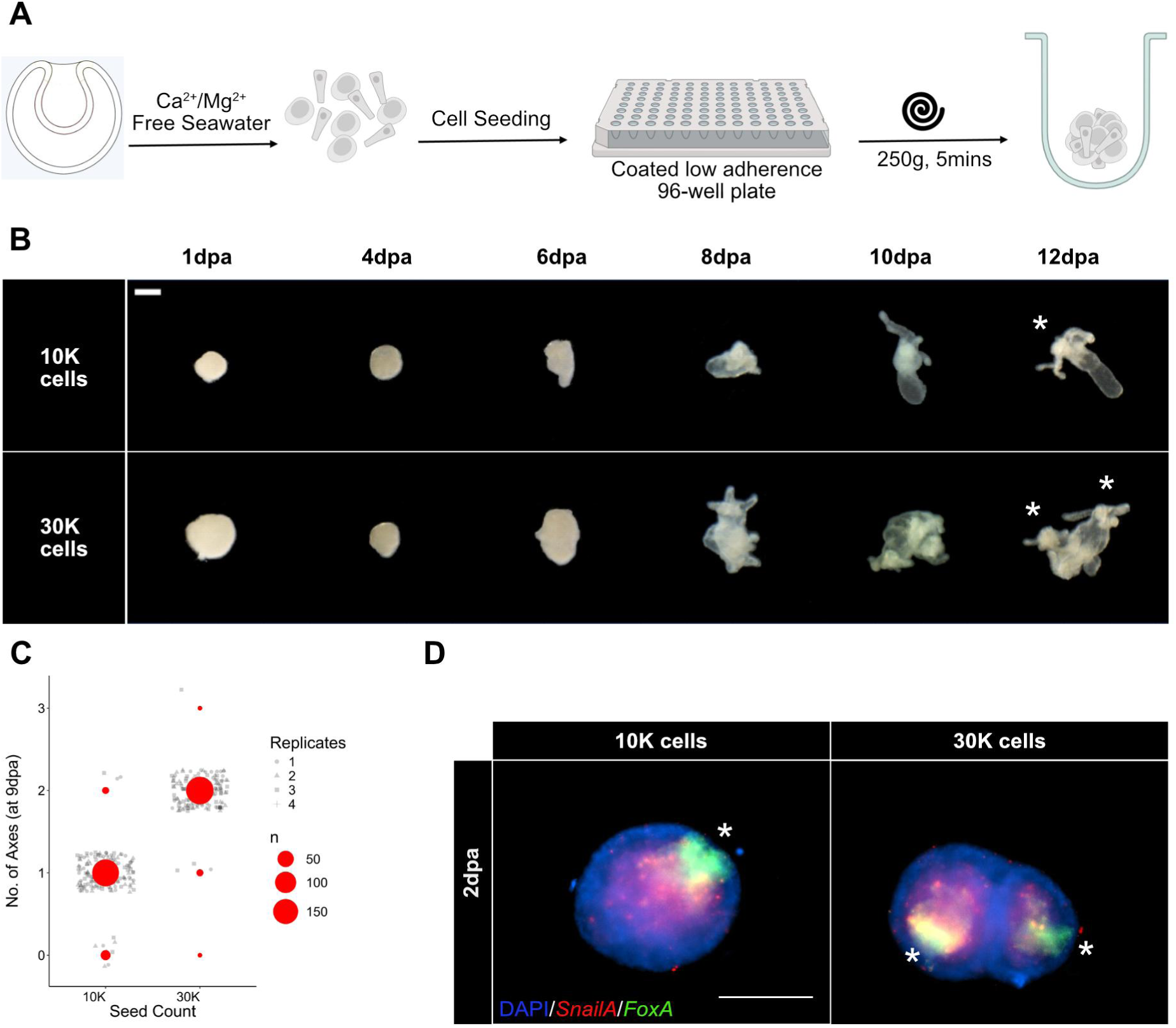
A standardized and scalable system for cnidarian gastruloid generation. **(A)** Schematic describing gastruloid generation from dissociated single cells of 24 hpf embryos. **(B)** Scaling of axes with increasing seed count in developing gastruloids.(Scale bar: 200μm). **(C)** Quantification of no. of axes across multiple biological replicates.10,000 cells robustly give rise to gastruloids with a single axis, while 30.000 cells generate 2 body axes.Jitter added to discrete values for complete sampling to be visualized. **(D)** Double fluorescent in situ hybridization (dFISH) staining for endoderm (*FoxA*+) and mesoderm (*SnailA*+) markers showing corresponding molecular scaling of axes. Asterisk denotes oral pole. dpa; days post-aggregation (Scale bar:100μm)

Having established a scalable and reproducible system to generate a single-axis polyp, we wished to monitor the morphogenetic process of germ layer formation. To follow the epithelization of the ectodermal cells, the aboral ectoderm of early gastrulae were dissected and dissociated from sfGFP-𝛃-catenin knock-in embryos (Lebedeva et al. 2025) followed by aggregation and imaging. As the ectodermal cells re-establish cell junctions, 𝛃-catenin is recruited to the apical-lateral membrane of all cells. Ectodermal epithelialization started as islands of a few cells, which then expanded to nearly complete epithelialization by 1 dpa **(Fig. 2A-F’)**. Additionally, the ectodermal cells formed 𝛃-catenin+ membrane junctions with neighboring ectodermal cells through a previously described mode of short range sorting **(Supplementary Video 1)**, as with ectodermal cells of *Hydra* (Hobmayer et al., 2016). Recent work also demonstrated the role of the lateral polarity gene product Lethal giant larvae (lgl) and NCAM2 in establishing proper epithelialization and apico-basal polarization of *Nematostella* ectodermal cells (Atajanova et al., 2024; Postnikova et al., 2025).

**Figure 2.**
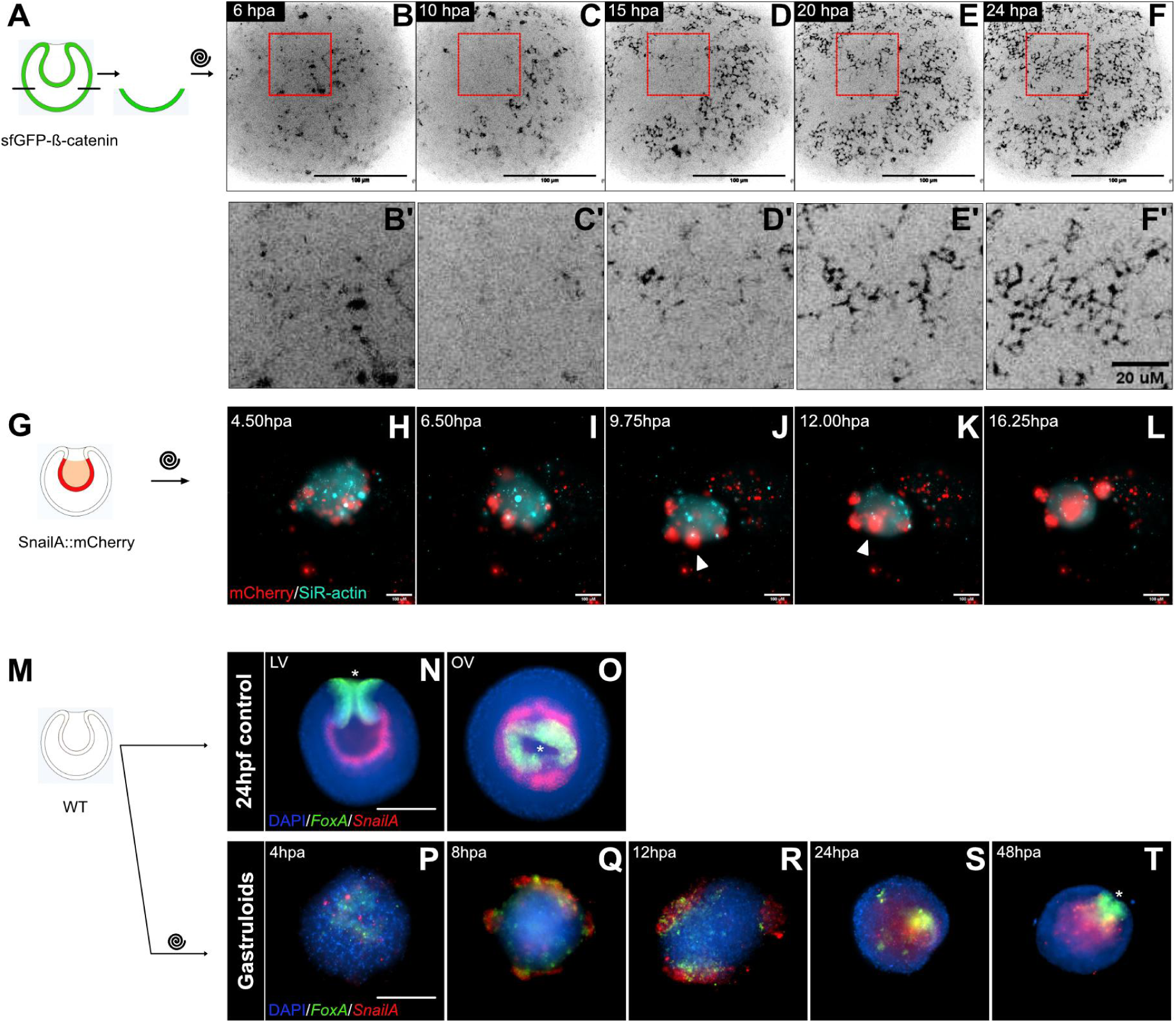
Distinct morphogenetic germ layer behaviors in *Nematostella* gastruloids. (A-F’) Rapid epithelialization of aboral ectodermal cell aggregates within a day, visualized using superfolderGFP-β-catenin transgenic line. Red square (B) represents same patch of cells for closeup across all timepoints (B’-F’). **(G-L)** Ingression of peripheral mesodermal clusters visualized by *SnailA::mCherry* (red). SiR-actin (blue) was used to counterstain all cells. White arrow represents points of ingression of the mesodermal cluster. **(M-T)** dFISH of *foxA* (green) and *snailA* (red) marking endodermal and mesodermal expression domains, respectively, in undissociated mid-gastrulae (N-O), and 4h, 8h, 12h, 24h and 48h post aggregation (hpa) (P-T). LV and OV stand for Lateral and Oral view, respectively. Asterisk denotes oral pole. DAPI (blue) was used as a counterstain of all cells. Twister symbol represents dissociation and reaggregation of cells (Scale bar:100μm)

The morphogenetic movement of the mesoderm was visualized by dissociation and reaggregation of *SnailA::mCherry* transgenic line embryos. *SnailA* is a marker of the whole inner cell layer, which has mesoderm-like profiles (Steinmetz et al. 2017; Haillot et al. 2025), including the expression of Cadherin1 upon downregulation of the ectodermal Cadherin3 during the Cadherin switch (Pukhlyakova et al., 2019). Surprisingly, instead of forming the inner cell layer from the start, the initially randomly distributed mesodermal cells (mCherry+) first formed cell clusters with other mesodermal cells at the periphery of the aggregate, before ingressing collectively as a cluster **(Fig. 2K)** to form the inner layer **(Fig. 2G-L)**. This is the behavior of all the mesodermal cells in the aggregate, including those located initially in the gastruloid center, previously thought to remain there through gastruloid development. Thus, the initially randomly distributed individual mesodermal cells either actively migrate or passively are pushed out to the periphery due to differences in cell adhesion forces in comparison to the ectodermal cells. Notably, at the surface, they tend to attach to other mesodermal cells forming rather compact clusters of about 20 cells **(Supplementary Video 2)**. On average, each gastruloid has about 3 clusters attached prior to ingression at 12hpa (hours post aggregation) **(Fig. S2.1)**. Notably, usually one of the clusters suddenly ingresses between 10-16 hours post aggregation, presumably at regions that have not yet completed epithelization **(Fig. S2.3)**.

In order to describe the morphogenetic behavior of the endodermal cells, marked by the conserved endoderm transcription factor gene *foxA*, we used fine temporal dFISH **(Fig. 2P-T)**. The results showed that an initially randomly distributed *foxA*+ endoderm **(Fig. 2P)** attached and intercalated with the *snailA*+ mesodermal cells already from 8hpa. This, together with the fact that mesodermal cells form their clusters by migration and differential adhesion, is suggestive that the endodermal cell population is also initially migratory post-dissociation. As the mesodermal cluster ingressed into the gastruloid, the multiple endodermal foci visible at around 12hpa **(Fig. 2R)** coalesced into a single primary pole. At 24hpa, multiple smaller poles were still visible **(Fig. 2S)** suggesting some underlying possible mechanism of cluster competition to choose a primary winning pole. While *foxA*+ cells exhibited intercalation, *brachyury*+ cells which are also primarily endodermal, clustered similarly with the mesoderm like *foxA*+ cells but with a clear boundary and no signs of intercalation **(Fig. S2.2)**. While the OA axis appears to be reestablished by 24hpa **(Fig. 2S)**, the endodermal cells at this point were still a clump of cells. Endoderm epithelialization and invagination was complete by 48hpa **(Fig. 2T)** thus, reestablishing the initial body plan **(Fig. 2N)**. This body plan is roughly comparable with the postgastrula planula larva. Thus, there is only about one day of delay in body axis establishment between gastruloid and normal embryo.

Notch signaling has been recently shown to induce the endoderm at the border of *Delta*+ (ligand) mesoderm and *Notch*+ (endoderm) (Haillot et al. 2025), making it a promising candidate regulating the endoderm-mesoderm boundary in *Nematostella*. Given that the endodermal cells intercalate within mesodermal clusters **(Fig. 2Q,R)**, we first checked if disrupting Notch signaling modulated this boundary in the gastruloids. Notch signaling inhibition with LY-411575 caused disruption of the endoderm-mesoderm boundary **(Fig. S3.5)** followed by subsequent depolarization of *foxA*+ endoderm in gastruloids observed at 2dpa **(Fig. 3A)**. The resulting gastruloids also lost their ability to make oral structures such as a pharynx and tentacles, in a LY-411575 concentration dependent manner **(Fig. S3.1)**. Interestingly, a similar depolarization was also observed in BMP morphant gastruloids implying potential cross talk of multiple pathways in regulating endodermal polarity (Kirillova et al. 2018). Since ectopic activation of Notch signaling could induce ectopic endoderm in early stage embryos (Haillot et al., 2025), we asked whether the stained endodermal population were the initial pre-dissociation endodermal cells, or if they arose newly through reinduction during self-organization. Previous work showed that the presence of the endodermal cells, which also express the organizer genes *wnt1* and *wnt3* in the gastrula prior to dissociation, is essential to confer self organizing potential to the gastruloids (Kirillova et al. 2018). However, this experiment was carried out by dissecting the blastopore lip, which could have also removed other essential cell types in the ectoderm and mesoderm. Since Notch signaling inhibitor LY-411575 specifically ablates the endodermal cell state without affecting ectodermal or mesodermal identity (Haillot et al. 2025), we sought to find out if pre-existing endoderm is essential. Confirming the results of the previous study, complete ablation of the organizer cells expressing *wnt1/wnt3* in the embryos **(Fig. S3.2A)** led to lack of axis establishment in the gastruloids **(Fig. S3.2B)**. While this experiment does not rule out induction of new cells, it suggests that the initial endodermal cells are essential to reestablish the axis.

**Figure 3.**
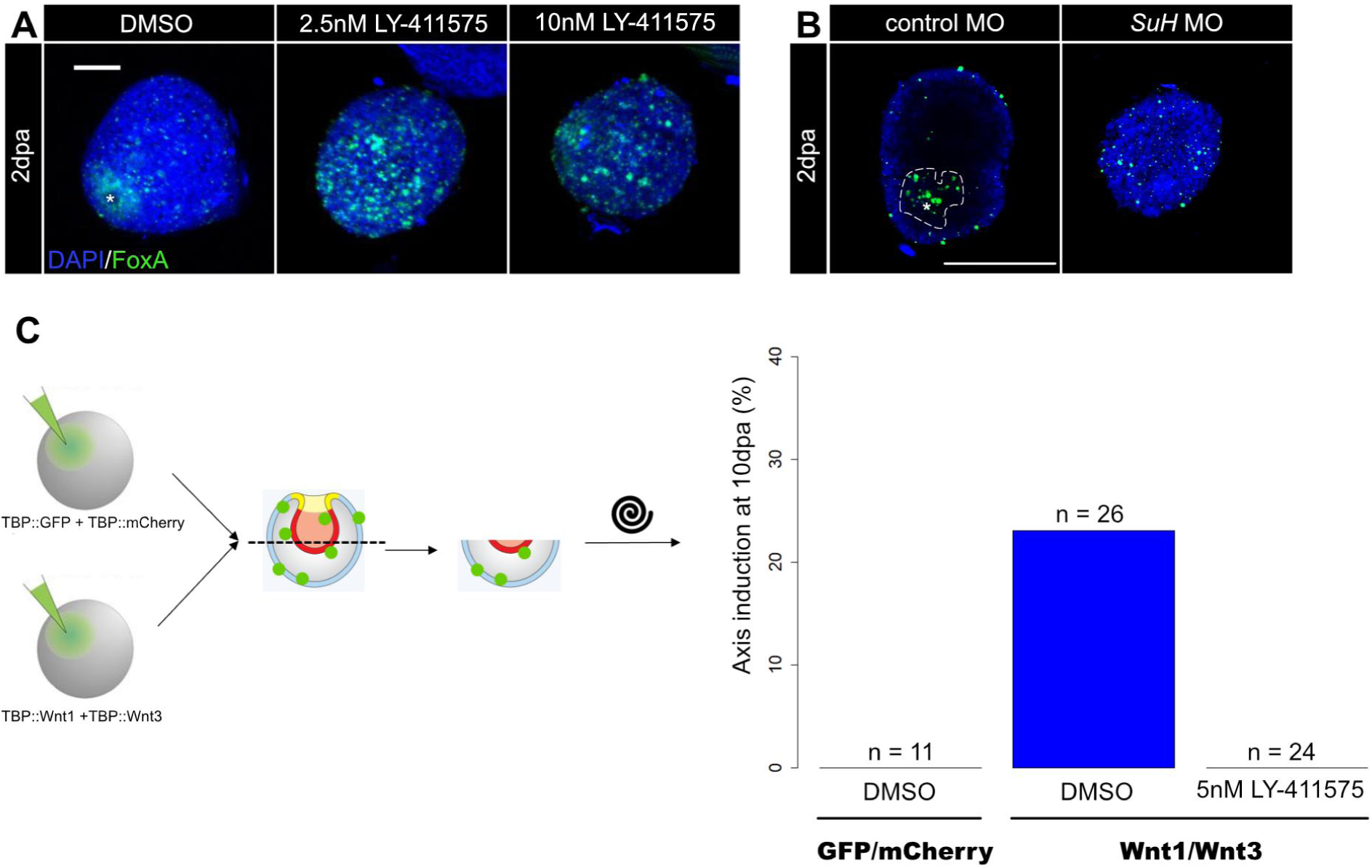
Notch signaling maintains endoderm polarity. **(A)** Gastruloids treated for 2 dpa with either DMSO or Notch signaling inhibitor LY-411575 and stained for *FoxA* depict dose dependent loss of endodermal polarity. Surface intensity profiles depicting the same. Asterisk denotes oral pole. dpa;days post-aggregation. (Scale bar: 50μm). **(B)** Gastruloids made from *SuH* Morpholino injected embryos phenocopy LY-411575 treatment compared to control Morpholino. Dotted lines denote polarized endoderm in control. (Scale bar: 100μm). **(C)** Primary axis induction (10 dpa) in gastruloids made from embryos co-injected with head organizers Wnt1/Wnt3 (green dots represent ectopic co-expression of constructs) and treated with either DMSO or LY-411575 shows Wnt signaling mediated gastruloid axis formation is Notch-dependent.

To understand which components of the Notch signaling cascade regulate this border, we first characterized an expanded panel of annotated canonical Notch signaling components in embryos **(Fig. S3.3A)**. The primary axis is reestablished in the gastruloids by 24hpa **(Fig. 2P-S)**. This coincides developmentally with the transition from 24hpf to 48hpf (hours post fertilization) embryos and therefore, we decided to validate the components in these embryonic stages. Inhibiting Notch signaling by LY-411575 treatment led to the downregulation of both the *notch* receptor and its ligands *delta* and *DLL*. This suggests that Notch signaling positively regulates itself **(Fig. S3.3B)**. The transition between 24 and 48hpf also exhibits a split of the expression domains of *delta* and *DLL* within the mesoderm. *Delta* has an expression maximum at the boundary with the endoderm, whereas *DLL,* a paralog of *delta,* is expressed interestingly in the aboral-most mesoderm, at the boundary with the apical tuft **(Fig. S3.3A)**. As the Notch receptor is expressed also in the aboral ectoderm, this receptor and ligand boundary might possibly play other roles, such as in the development of the apical tuft. Finally, the expression domain of annotated Notch signaling coactivator gene *Suppressor of Hairless* (*SuH*), also matched that of *notch* receptor. SuH was reported to be a potential Notch coactivator in *Nematostella* cnidogenesis (Marlow et al. 2012). Knockdown of *SuH* was also reported to phenocopy the Notch signaling inhibition phenotype in early embryos. Therefore, we also wanted to check if *SuH* was also involved in maintaining polarity of the endoderm in gastruloids. Our results show that gastruloids made from *SuH* MO injected embryos also phenocopy LY-411575 treated gastruloids **(Fig3B)**, thus suggesting involvement of SuH and the canonical Notch signaling cascade in endoderm polarity maintenance.

Finally, given the role of Notch signaling in maintaining endodermal polarity, we also wanted to see if there was any interaction between Notch and Wnt signaling in the gastruloids. Gastruloids made from only aboral ectodermal cells are incapable of generating a body axis and germ layers, while the addition of aboral cells co-injected with *wnt1/3* are can rescue this phenotype and will reestablish the OA axis (Kirillova et al. 2018). To test if there was any interaction between the Notch and Wnt signaling pathways, we repeated this experiment with the gastruloids grown in Notch signaling inhibitor LY-411575 instead. We confirmed that the addition of wnt1/3 expressing cells to the aggregates from aboral ectoderm rescued the axis forming capacity of the aggregates (Fig. 3C). However, when the Notch signaling inhibitor LY-411575 was added, the ability to generate an axis was lost completely, despite the presence of *wnt1/3* expressing cells **(Fig. 3C).** This suggests that Notch signaling acts downstream of Wnt Signaling in axial identity establishment. Since *notch* and *delta* expression in the early embryo were recently shown to depend on Wnt signaling (Haillot et al. 2025), our results demonstrate the presence of a feedback loop between Notch and Wnt signaling, which is essential for organizer symmetry breaking in the gastruloids.

To unravel how Notch signaling modulated the identity of the endoderm post-gastrulation, we first checked if it continues to play the role of maintaining endodermal gene expression as previously reported (Haillot et al. 2025). For this, we used embryos treated with LY-411575 from 24hpf to 48hpf (post-gastrulation time window) and stained for key endodermal markers *foxA* and *brachyury* as well as 𝛃-catenin target *axin* (Kraus et al. 2016; Lebedeva et al. 2025). Notch signaling inhibition led to the downregulation of all these genes **(Fig. S4.1)** showing that it still continues to play a role in maintaining endodermal identity. To identify the broader contribution of Notch signaling in modulating the identity of the endoderm and the other germ layers, we generated scRNAseq datasets of the above mentioned LY-411575 treated (and DMSO control) embryos where cluster identities were annotated by label transfer from a reference atlas (Cole et al. 2024) (detailed in Methods). As validation of sufficient drug treatment strength, we found many of the genes previously reported to be downregulated by Notch signaling inhibition to also be downregulated in our datasets (**Fig. S4.3**). Upon cluster annotation, we could observe an expansion of the *soxC*+ secretory progenitor cells as well as a truncation of cnidocyte maturation, with a lack of *soxA*+ mature cnidocytes in the LY-411575 treated embryos **(Fig. 4A)**. This is in alignment with earlier works that have demonstrated the dual role of Notch signaling in inhibiting neurogenesis by limiting the pool of progenitors, as well as promoting cnidogenesis (Richards and Rentzsch, 2015; Marlow et al. 2012; Plesseir and Marlow, 2025). We carried out Weighted Gene Co-expression Network Analysis (WGCNA) analysis to identify gene modules associated with specific cell type responses to the drug treatment **(Fig. S4.4).** One such interesting module was the module specific to the endomesoderm, here termed as the pink module. This module included genes whose expression were primarily mesoderm-specific in control embryos, however, expanded into the endoderm in the LY-411575 treated embryos **(Fig. 4C)**. This indicated that Notch signaling inhibition did not only lead to a downregulation of endodermal identity as previously seen **(Fig. S4.1)**, but also to an adoption of mesodermal features, hence defining a transient endomesodermal state **(Fig. 4A,B)**. Among the expanding genes of the pink module, an interesting candidate we investigated further was *Cadherin1* **(Fig. 4C)**.

**Figure 4.**
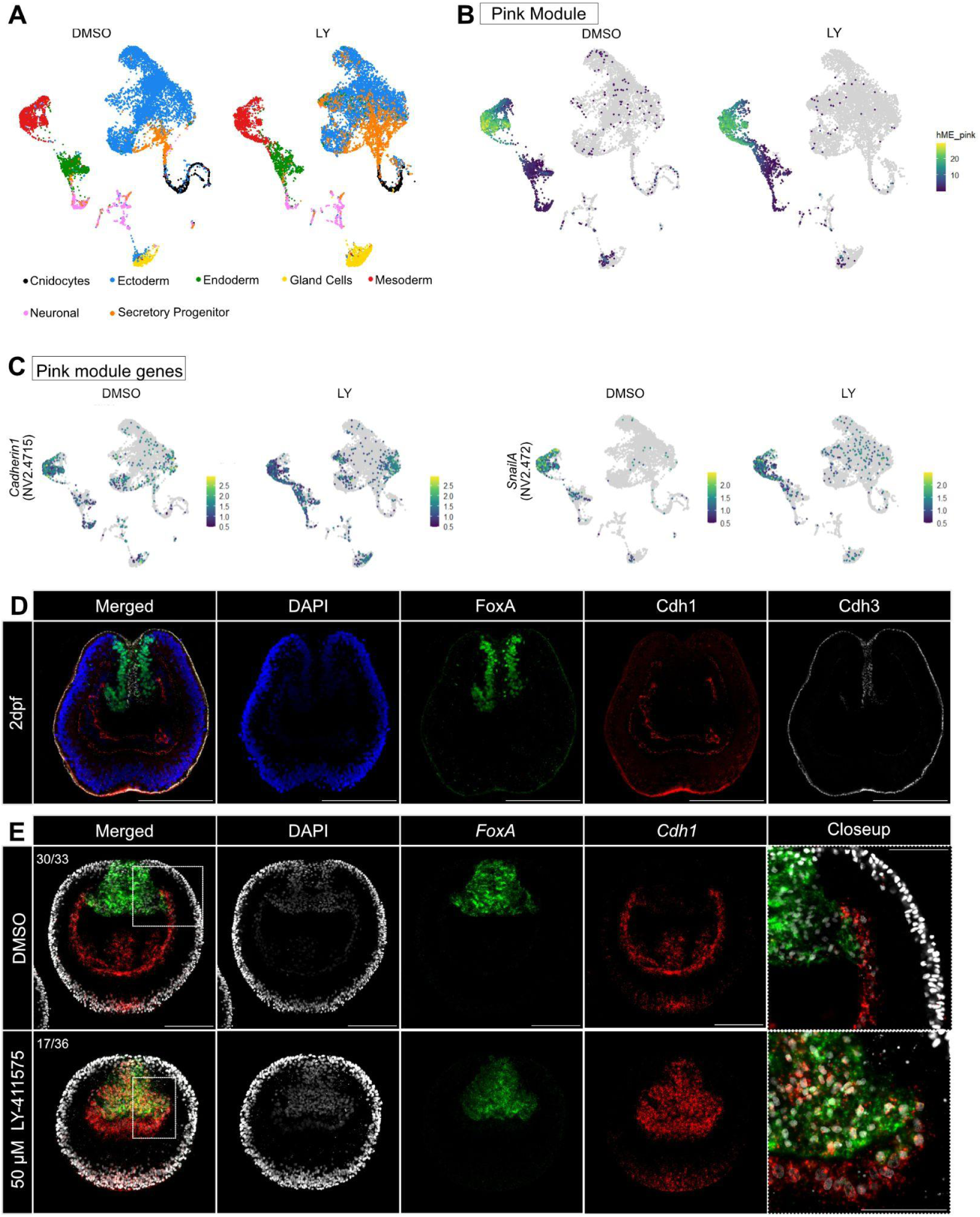
Notch signaling inhibition converts endoderm to endomesoderm. **(A)** UMAP dimensional reduction of reference annotated scRNAseq cell clusters comparing DMSO and LY-411575 drug treated embryos, treated in a window of 24 to 48hpf. **(B)** Feature plot highlighting pink module eigengene expression across both DMSO and LY-411575 treated embryos. **(C)** Feature plot of key pink module genes, *cdh1* and *snailA*, highlighting expansion of mesodermal genes into endoderm upon Notch signaling inhibition. **(D)** Triple antibody staining in 2dpf embryos for FoxA, Cdh1 and Cdh3 shows the endoderm expressing Cdh3 while the mesoderm expresses Cdh1 after the Cadherin switch. (Scale bar: 100μm). **(E)** Notch signaling inhibition causes loss of Cadherin switch with endoderm co-expressing *foxA* and *cdh1* (Scale bar: 100μm). Closeups depict the boundary between endoderm and mesoderm. Note the sharp, mutually exclusive boundary between *foxA* and *cdh1* in the DMSO control, compared to the overlapping expression in LY-411575 treated embryos (Scale bar: 50μm).

During gastrulation, *Nematostella* the early ubiquitous blastodermal Cadherin3 (Cdh3) becomes downregulated and replaced by Cadherin1 (Cdh1) in the invaginating mesoderm. As a result of this Cadherin switch, Cdh1 and Cdh3 form a sharp meso-endodermal boundary (Pukhlyakova et al. 2019) **(Fig. 4D)**. Cdh3 was previously shown to play an important role in epithelialization of the ectoderm in *Nematostella* aggregates (Pukhlyakova et al. 2019). To test whether Cdh1 plays a role in adhesion of the mesoderm, we first knocked down *cdh1* expression by injecting *cdh1* morpholinos (MO) in embryos of a mesoderm-specific reporter line (𝛃*-Laminin::eGFP-CAAX)*, which we generated in order to monitor the organization of the mesoderm **(Fig. S4.5A)**. Upon *cdh1* knockdown (KD), the mesoderm loses its epithelial architecture in the embryos **(Fig. S4.5B)** in line with Cdh1 expected role in mesoderm adhesion. However, since Cdh1 is also expressed in the aboral ectoderm at the planula stage (Pukhlyakova et al., 2019), *cdh1* knockdown also appeared to disrupt the ectoderm, which could also indirectly affect mesoderm organization. Therefore, to show mesodermal specificity, we knocked down 𝛃*-catenin,* which converts the identity of the whole embryo into embryonic mesoderm (Haillot et al., 2025), including an ectopic expression of mesodermal markers *snailA* and *cdh1* throughout the embryo (**Fig. S4.5C**). These mesodermalized 𝛃-catenin KD embryos were injected either with control MO or with *cdh1* MO, and then proceeded to dissociate and reaggregate these embryos. We observed that gastruloids made from 𝛃*-catenin*/control KD embryos aggregated properly, however, 𝛃*-catenin/cdh1* KD aggregates fell apart, confirming the function of Cdh1 in adhesion of mesodermal epithelial cells **(Fig. S4.5C)**.

To verify the loss of Cadherin switch observed in the LY-411575 treated embryos in the scRNAseq dataset, we used dFISH which revealed a disruption of the mutually exclusive endodermal and mesodermal domains leading to the expected expansion of the mesodermal *cdh1* expression into the *FoxA*+ endoderm **(Fig. 4D)**. Additionally, we also observed a strong global downregulation of *cadherin3* expression **(Fig. S4.2)**, potentially through Notch signaling mediated downregulation of Wnt signaling **(Fig. S4.1)**.

To further validate this, we monitored changes of cell states throughout development of the gastruloid using single cell RNAseq. We used a 24hpf gastrula library as our T0 control timepoint **(Fig. 5A)**. After clustering, we annotated the clusters by query mapping to the latest *Nematostella vectensis* scRNAseq atlas (Cole et al., 2024). We started by checking the expression of previously identified pink module genes, *cdh1* and *snailA* **(Fig. 4C)**. Indeed, both *cdh1* and *snailA* are upregulated in the endodermal cluster during early stages of aggregation, confirming an early transient endomesodermal cell state. Subsequently, *cdh1* gets down-regulated by 24hpa and 48hpa in this cell cluster adopting a more typical endodermal cell state **(Fig. 5B)**. This coincides with the reestablishment of cell contacts between the endoderm and mesoderm followed by proper resegregation of endoderm and mesoderm. The presence of the transient endomesodermal cell states marked by *foxA*/*cdh1* expression was also verified by Hybridization Chain Reaction (HCR) with an initial major overlap between *foxA*+ and *cdh1*+ domains at 4 hpa and a receding overlap with continued development at 24 hpa **(Fig. 5C)**. In the transcriptome of the transient endo-mesodermal cell state, we found many genes characteristic of undergoing epithelial mesenchymal transition (EMT), which is consistent with the previous finding that the mesodermal cells undergo a partial EMT phenotype during gastrulation (Kraus and Technau, 2006).

**Figure 5.**
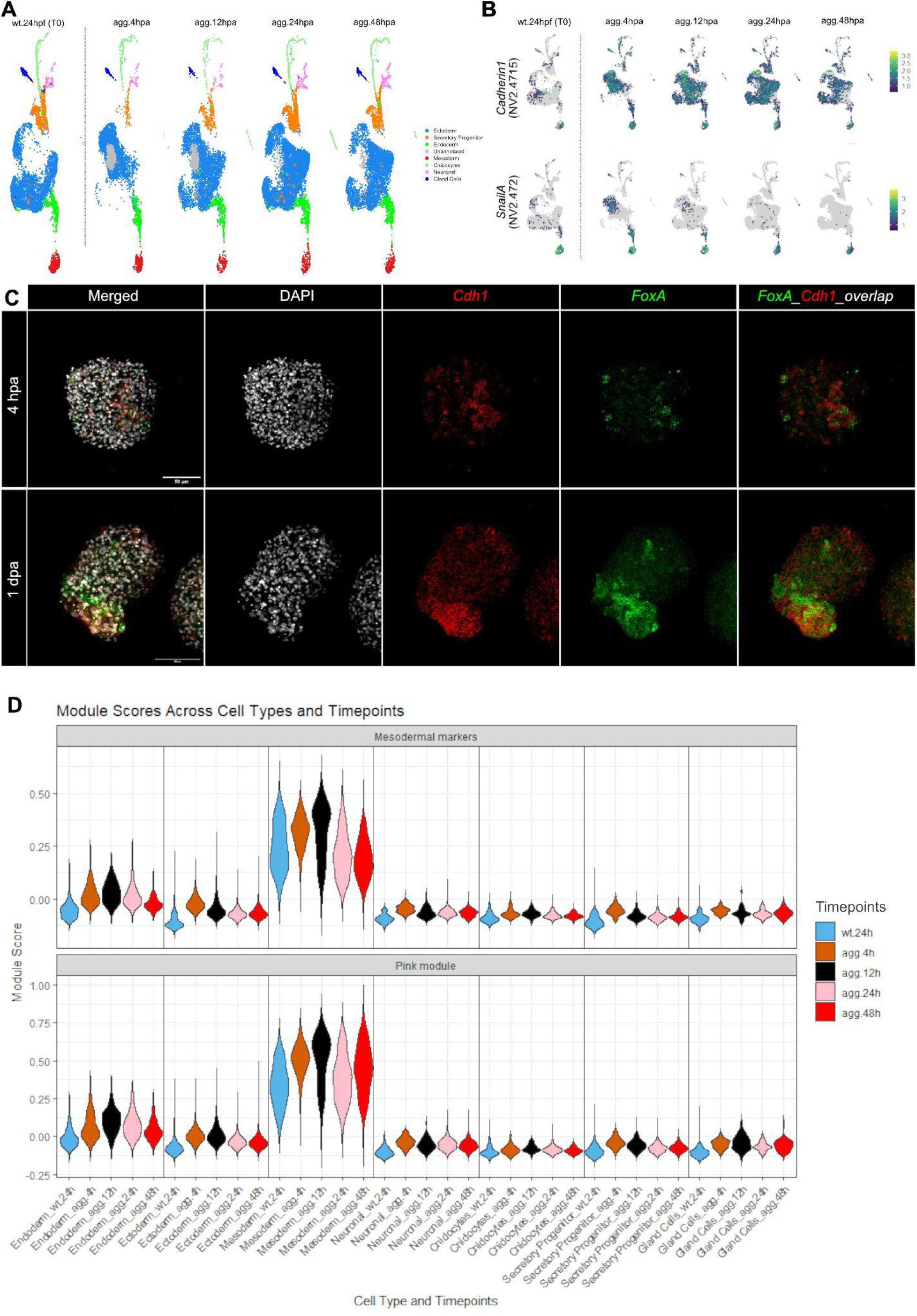
scRNA-seq gastruloid atlas confirms endoderm conversion to endomesoderm. **(A)** UMAP dimensional reduction showing different tissue cell clusters from the gastruloid time course starting from undissociated 24 hpf gastrula (T0) to dissociated and reaggregated gastruloids from 4 h to 48 hpa. **(B)** Feature plot showing expression dynamics of example pink module genes *cdh1* and *snailA* across gastruloid development time course. Note the transient expansion of mesodermal markers into the ecto-endodermal territory. **(C)** HCR expression profile of endodermal marker *foxA* and mesodermal marker *cdh1* at 4 hpa and 24 hpa in gastruloid development. **(D)** Pink module and mesodermal module projection scores calculated for all annotated clusters across timepoints of gastruloid development.

We also calculated the module score of the entire pink module gene set for our gastruloid development dataset **(Fig. 5D)**. This score measures the average expression of the genes in the pink module, adjusted by subtracting the background expression of control gene sets that have similar expression levels. As expected, the score is highest for the mesodermal cluster as the expression of the genes are generally mesoderm specific, however, we also observed an elevation in the endoderm cluster especially in the gastruloids compared to the pre-dissociation 24hpf embryo **(Fig. 5D).** This was also evident in the module score ranking with the gastruloid endoderm having the second highest rank after the mesoderm **(Table S3)**. To show similarity of the pink module gene set to the mesoderm, we also identified differentially expressed genes specific to the mesodermal cluster from the wildtype scRNAseq atlas (Cole et al., 2024) and calculated the module score again for these genes in the gastruloid dataset. In alignment with the pink module, scores for the mesoderm module also behaved in a similar fashion with highest scores in the mesoderm and an elevation in the endodermal clusters of the gastruloid libraries **(Fig. 5D)**. Taken together this confirms the transient conversion of the endoderm to an endomesodermal cell state after dissociation and following reaggregation.

## Discussion

Our findings reveal a unique mechanism of self-organization where both germ layer topology and axial polarity are reestablished through coordinated Notch and Wnt signaling. In normal embryos, Notch signaling plays a crucial role in segregating the endoderm and mesoderm identity, by inducing and maintaining the endodermal identity (Haillot et al., 2025), which is marked by the sharp boundary between ecto/endodermal Cadherin3 and mesodermal Cadherin1 (Pukhlyakova et al., 2019). Upon dissociation, the interaction of the mesodermal Delta ligand with the ecto/endodermal Notch receptor and hence Notch signaling in the endodermal tissue becomes disrupted (**Fig. 6**). This abolishes transiently the maintenance of the endodermal identity signaling inhibition, hence leading to an endo-mesodermal profile. In embryos, this is mimicked by treatment with the Notch signaling inhibitor LY-411575, which leads to an expansion of (mesodermal) Cdh1 expression into the endodermal cells. In gastruloids, the expanded Cdh1 expression in single FoxA+ endodermal cells presumably allows these endomesodermal cells to adhere to the Cdh1/SnailA+ mesodermal cells during the process of axial polarization and restoration of the correct germ layer identity topology **(Fig. 5).** This also explains the observed clustering of these endo-mesodermal cells with mesodermal cells **(Fig. 2P-T)** prior to reestablishing endodermal identity through reinitiation of Notch signaling. Moreover, the transcriptomic profile suggests that due to the dissociation, the cells also adopt transiently a partial EMT-state, presumably facilitating the migratory behavior prior to the clustering. This is in line with the partial EMT phenotype of mesodermal cells during normal gastrulation (Kraus and Technau, 2006). In summary, while Notch signaling is required for the axial polarization, its transient disruption due to the dissociation is necessary to allow the attachment of endodermal and mesodermal cells. If Notch signaling is continuously inhibited in aggregates by treatment with LY-411575, reclustering of endodermal and mesodermal cells and hence axial polarization is abolished.

**Figure 6.**
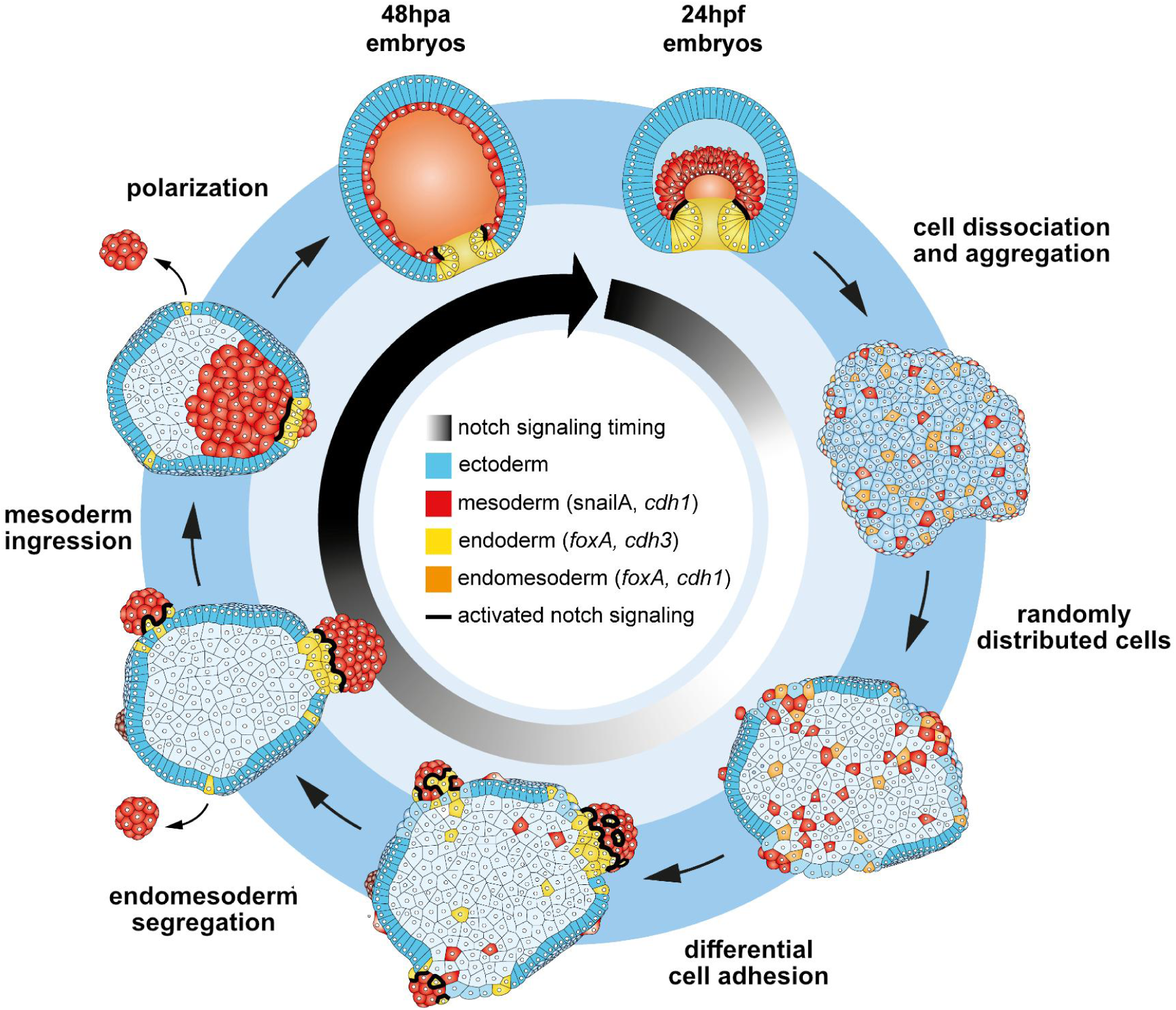
Summary of self-organization of *Nematostella* gastruloids. See text for explanation.

Endoderm undergoing EMT is a behavior that can also be found in some mammals such as mouse embryos where the definitive endoderm undergoes EMT to leave the primitive streak along the mesodermal wings before undergoing MET to form the gut endoderm (Nowotschin et al. 2019).

While upstream signaling pathways driving mouse epiblast migration are well studied, it is still unclear what drives the EMT-MET cycle of the definitive endoderm (Zorn and Wells, 2009). Based on our results, cell-cell signaling pathways like Notch signaling could be a plausible candidate. Interestingly, in zebrafish embryos, endoderm internalization is mediated by activation of the N-Cadherin (Cdh2), triggering them to become migratory (Giger and David, 2017). Despite not being orthologs, both *Nematostella* and bilaterian cadherins exhibit functional convergence in mediating cell adhesion and actin binding dynamics (Clarke et al. 2016). In addition, their cadherin switch from Cdh3 to mesoderm specific Cdh1 during gastrulation (Pukhlyakova et al. 2019), parallels E- to N-Cadherin switch in vertebrates (Mayran et al. 2023). It remains to be seen if the *cdh1* expression in the endoderm of the developing *Nematostella* gastruloids can trigger endoderm EMT.

Our results also have important implications on the theory of self-organization. Turing-type reaction-diffusion models have provided a theoretical framework for pattern formation and self-organisation from almost homogenous conditions. In these classical Turing models, the short-range autocatalytic activator would cross-activate a long-range inhibitor, leading to local activation and lateral inhibition (Turing, 1952, Gierer and Meinhardt, 1972). Such a system is able to give rise to either single (body) axis or to periodic patterns as observed in cases like digit patterning.Wnt/𝛃-catenin signaling with its slow and short diffusing ligands and quick intracellular 𝛃-catenin stabilization is a strong candidate for a Turing-type activator capable of generating polarity from initially homogeneous conditions (Pond et al. 2020; Turing, 1952). Recent evidence from mammalian gastruloids demonstrate that Wnt polarity can emerge from an initially homogenous population of cells through self-reinforcing 𝛃-catenin/Tcf feedback loops (McNamara et al. 2023). Pattern formation by a Turing-type reaction-diffusion system assumes that self-organization is based solely on the re-establishment of signaling gradients, without the requirement of cell sorting.

In mammalian systems, the number and size of poles are regulated by multiple factors such as size and shape of gastruloids (Moris et al. 2020), adhesion based sorting (Turner et al. 2017; McNamara et al. 2023), as well as production of Wnt antagonists such as Dkk1 (Dickkopf-1) (Belo et al. 1997; Glinka et al. 1998; Kawano & Kypta, 2003; Kemp et al. 2005), which meet the key criteria of a Turing inhibitor. In the cnidarian *Hydra,* the head is maintained by an autoregulatory Wnt3/𝛃-catenin circuit (Nakamura et al. 2011), with the number of heads controlled by a Turing-type inhibitor Sp5, whose knockdown leads to the expected multiheaded polyp phenotype, as well as multiple head organizer poles in aggregates made from dissociated and reaggregated polyps (Vogg et al. 2019; Galliot and Wenger, 2025; Narayanaswamy and Technau, 2025). In *Nematostella* gastruloids, it was previously shown that the presence of some *wnt1/wnt3* expressing cells is required to establish a body axis (Kirillova et al. 2018). These *wnt1* and *wnt3* expressing cells are found in the blastopore lip of the gastrula (Kusserow et al. 2005), which also can act as axis organizer when transplanted to an aboral position (Kraus et al. 2016). They form a feedback loop with *foxA* and *brachyury* (Schwaiger et al. 2022) and together they demarcate the tissue with transcriptomic profile of the bilaterian endoderm and hence is here termed endoderm (Steinmetz et al. 2017; Haillot et al. 2025). Notably, self-organisation of a whole sea anemone polyp from gastruloids does not only imply the formation of a body axis, but also the re-establishment of the topology of three germ layer identities. Therefore, we propose a dual, connected activator system with Wnt/𝛃-catenin signaling driving broad axial patterning and Notch signaling as its coactivator in establishing and maintaining endodermal identity.

We emphasize the hitherto unappreciated significance of Notch signaling for the process of self-organization in sea anemone gastruloids. Of note, recent work in regenerating mouse intestinal epithelium demonstrated that Notch/Delta-dependent lateral inhibition is coupled to a bistable cell fate switch (Schwayer et al. 2025). Here, transient tissue-density-dependent heterogeneity in the mechanosensor YAP1 pre-patterns the expression of the Notch ligand DLL1 via the FOXA1, and DLL1–Notch feedback creates two stable fate states (high-DLL1’sender’ cells and low-DLL1’receiver’ cells). This bistability allows patterns to remain even after the initial YAP1 heterogeneity is lost, providing a ‘memory’ of earlier symmetry-breaking signals. Perturbing Notch signalling alters both the fraction and spatial arrangement of DLL1+ cells, demonstrating that lateral inhibition here not only refines boundaries, but stabilizes the pattern formation once completed. In the sea anemone gastruloids reported here, due to the rapid epithelialization and small size there may not be a requirement for classical fast diffusing Turing inhibitors. This system could achieve axis polarization simply through Wnt/Notch signaling driven symmetry breaking, Notch mediated boundary refinement and potential cadherin dependent cell sorting. By analogy with the mouse intestinal regeneration model, such a Wnt/Notch circuit in gastruloids could operate via a similar bistable lateral inhibition module, preserving organizer versus non-organizer identity and stabilizing against later noise or fluctuations. This pattern scales with the number of cells, as 30,000 cell gastruloids robustly form polyps with 2 axes **(Fig. 1B,D)**. This shows that multiple organizer poles can emerge if they are at sufficient distance from each other, indicating the range of inhibition. This scenario mirrors reports of multiple axes in large mouse gastruloids (Ishihara and Tanaka, 2018), with larger tissue sizes warranting inhibitors for wavelength control as in Turing patterns. More experiments studying larger gastruloids would be needed to garner insights into the relevance of Turing models in axial pattern formation.

Cnidarians in general exhibit extreme regenerative capabilities with multiple species that can self-organize from being dissociated and reaggregated (Kirillova and Kraus, 2018). The various parameters and mechanisms underlying their self-organization have however only been studied in detail primarily in *Hydra* (Technau and Holstein, 1992; Technau et al. 2000; for review see Narayanaswamy and Technau, 2025) and *Nematostella* (Kirillova et al. 2018) and more recently in *Hydractinia* (Curantz et al. 2025). In *Hydra*, germ layer topology and axial polarity are largely decoupled with head organizers capable of emerging *de novo* driven by a combination of biochemical and mechanical cues (Narayanaswamy and Technau, 2025). In aggregates made from *Hydractinia* polyps, self-organization is a 2-step process with the first step being Sphingosine dependent i-cell migration towards the epidermis to form clusters, followed by Wnt signaling within these clusters that reestablish the head-foot axis. Additionally, all somatic cells are replenished by the i-cells (Plickert et al. 2012; Curantz et al. 2025). In *Nematostella* gastruloids, although not directly comparable in terms of ontogeny to *Hydra* or *Hydractinia*, exhibits an alternative mode of self-organization where axial polarity and germ layer organization are reestablished in parallel, mediated by Notch signaling.

The divergence of mechanisms across cnidarians indicate that the principles of self-organization among cnidarians may not be universal, motivating a need to describe the variable potential of developmental robustness observed within cnidarians (Narayanaswamy and Technau, 2025). Since the starting point are early embryonic cells, we consider the *Nematostella* gastruloids as a useful cnidarian *in vitro* embryonic self-organization system that is comparable to vertebrate organoids and gastruloids. We propose that the morphospace that is available to the organisms is larger than anticipated, reflecting a developmental robustness but which also may be a source for evolutionary plasticity of body plans on the basis of conserved regulatory networks. This comparison would help us learn more about the degree of developmental plasticity in early branching animals that may have allowed for the subsequent diversification of body plans.

## Materials and Methods

### Animal culture and spawning

*Nematostella vectensis* polyps were grown in 16%ₒ Nematostella medium (hereafter NM) at 18°C as previously described (Fritzenwanker et al., 2002). These animals were fed 5x a week with *Artemia* nauplii and spawning was induced by a 9h light exposure window at 25°C followed by incubation for 2.5 hours at 18°C. Eggs were then collected and fertilized with sperm for 45 mins. Fertilized eggs were dejellied in a solution of 3% L-cystein/NM solution (Genikovich and Technau, 2009) for 30 mins and then cleaned thoroughly to remove residual L-cystein using NM. Subsequently, embryos were then raised at 21°C.

### Gastruloid generation

Low adhesion 96-well plates were prepared by coating sterile non-coated 96-well plates with BIOFLOAT™FLEX coating solution (FaCellitate) using manufacturer instructions. Mid-gastrula stage embryos were then dissociated using Calcium Magnesium free seawater (hereafter CMF) until the suspension consisted of only single cells with no noticeable clumps (Kirillova et al., 2018). Tubes were filled up with filtered NM and centrifuged at 400 rcf for 4 mins. Supernatant was discarded and the cells were washed once with filtered PBS before being finally pelleted and resuspended in filtered PBS. The concentration and viability of the cells were then stained with ViaStain™AOPI staining solution (Nexcelom) and quantified using Cellometer X2 using parameters appropriate for gastrula stage cells. Resultant concentrations were used to calculate required volumes for different seed counts. 10,000 cells were seeded to generate standardized single axis gastruloids. The cells were then seeded in wells containing filtered NM and spun gently and quickly at 250g for 5mins. Plates were incubated at 21°C and cell debris were cleaned 1 dpa and gastruloid development was subsequently monitored.

### Live imaging

Gastruloid epithelialization (**Fig. 2A-F’**) was imaged by dissecting out the aboral ectodermal half of 𝛃-catenin-sfGFP KI line midgastrula embryos (Lebedeva et al. 2025) and dissociated and reaggregated as described previously. These gastruloids were allowed to develop until 3hpa when the cells had sufficiently adhered to each other, and then embedded in 2% low melting agarose and imaged on a spinning disk microscope. Site of ingression of mesodermal cells (*SnailA*+) (**Fig. S2.3**) were imaged by crossing 𝛃-catenin-sfGFP KI females and *SnailA::mCherry* reporter males and resulting embryos were used to generate gastruloids and were imaged similarly on the spinning disk confocal. Long term tracking of ingressing peripheral mesodermal (*SnailA*+) cell clusters during gastruloid development (**Fig. 2G-L**) was carried out free floating in NM within special teflon wells using the Light sheet microscopy (Viventis).

### Cloning and gene expression analysis (Colorimetric + Fluorescent)

Transcripts of genes of interest were amplified from cDNA using primers listed in **Table S1.** PCR products were cloned in pGEM-T (Promega, #A3600). Both FITC or DIG-labeled RNA probes were then generated through *in vitro* transcription with SP6 or T7 polymerase. In situ hybridization was carried out as described previously (Kraus et al. 2016; Haillot et al. 2025). After incubation with anti-Digoxigenin AP (1:4000) or anti-Fluorescein/Digoxigenin POD (1:100), embryos and gastruloids were washed 10 times 10 minutes with TBST (0.5% Tween). Background was removed by washing embryos 6 times with TBST (0.5% Tween) and kept overnight at 4°C. Double fluorescent in situ hybridization (dFISH) was performed similar to single in situ hybridization according to previously reported protocol (Tourniere et al. 2024). Fluorescent staining was revealed with the TSA Plus Fluorescein and TSA Plus Cy3 detection Kits (AKOYA Biosciences). Post staining, embryos were infiltrated with Vectashield (VectorLabs) and imaged.

Following modifications were made while staining gastruloids:

a. Embryos were grown in 0.5% DMSO from fertilization until prior to dissociation to improve staining in subsequently made gastruloids.
b. Given their much smaller volume due to compaction, gastruloids were regularly spun down between ISH steps to prevent loss of material.

Hybridization chain reaction (HCR) staining was done as previously reported (Choi et al. 2018) with some modifications. Probe pools for genes *cdh1* and *foxA* were generated (Molecular Instruments HCR v3.0) (**Table S2.**). Gastruloids were fixed at 4hpa and 1dpa respectively with 4% PFA diluted in 1xPBS for 1h at room temperature (RT), washed 3 times with PTw (PBS + 0.5% Tween-20) and stored in 100% Methanol at-20℃ until needed for staining. Gastruloids were washed with 50% Methanol/50% PTw 2 times followed by 3 washes in PTw. The rehydrated gastruloids were then washed 2 times for 5 min in 5xSSCT (5x Sodium saline citrate buffer, 0.1% Tween-20). They were then prehybridized in Probe hybridization buffer (Molecular Instruments) for 30 min at 37℃. They were then hybridized in hybridization buffer consisting of the probe set (0.8 pmol probe in 100 ul hybridization buffer) overnight at 37℃. Samples were then washed 4 times for 10 min and 2 times for 3 min with Probe wash buffer (Molecular Instruments) at 37℃, followed by 5 times 5 min wash with 5xSSCT at RT. Samples were subsequently incubated in Amplification buffer (Molecular Instruments) at RT for 30 min. Parallely, hairpins h1 and h2 (12pmol) were heated separately at 95℃ for 90s, snap cooled to RT in the dark, and added to 100ul of Amplification buffer. Gastruloids were then incubated in this hairpin/amplification buffer mix overnight at RT in the dark. Post signal amplification, samples were washed with 5xSSCT: one time for 1 min, three times for 5 min, and two times for 30 min. Samples were then infiltrated with Vectashield (VectorLabs) and imaged using the Leica Stellaris 5 CLSM.

### Inhibitor treatments

Inhibitor stocks were dissolved in DMSO. Notch signaling was inhibited using a specific gamma-secretase inhibitor (LY411575, MedChemExpress, #HY-50752). In the case of embryo treatments, a concentration range of 40μM-70μM was used, treating from 24hpf to 48hpf. For gastruloids, a much lower concentration range of 0.1nM-20nM was used due to their initial treatment as a clump of single cells, making them more sensitive to inhibition than compact embryos. The treatment window was from 0hpa to the time of sampling. To generate scRNAseq libraries to capture Notch inhibition response of embryos, 50μM LY411575 was used.

### Antibody staining

Staining was carried out as described previously (Pukhlyakova et al. 2019). In brief, samples were fixed in 4% PFA in 1xPBS for 1h at RT, following which they were incubated in ice cold acetone at-20℃ for 7 min. For samples being stained for cdh1 alone, samples were fixed in Lavdovsky’s fixative as reported previously (Pukhlyakova et al. 2019). Then they were washed 5 times in PTx (PBS + 0.2% Triton-X) and blocked in blocking solution (20% Sheep serum, 1% Bovine serum albumin (BSA) in PTx) for 2h at RT. Primary mouse anti-Cdh3 antibody (1:1000), rat anti-Cdh1 (1:500), rabbit anti-FoxA (1:500) and rabbit anti-SnailA (1:500) were diluted in blocking solution and samples were incubated in the relevant primary antibodies overnight at 4℃. They were then washed in PTx at RT (10×10 min each) and blocked again in blocking solution for 2h at RT. They were then incubated in relevant secondary antibody solutions of goat anti-mouse Alexa Fluor 633 (1:1000), goat anti-rat 568 (1:1000) and goat anti-rabbit 488 antibodies with DAPI (1:1000) diluted in blocking solution overnight at 4℃. They were then washed in PTx (10×10 min) and infiltrated with Vectashield (VectorLabs) and imaged using the Leica Stellaris 5 CLSM.

### Plasmid and morpholinos microinjection

For knockdown experiments, previously published antisense translation blocking Cadherin1 morpholino (Pukhlyakova et al. 2019), splice blocking SuH morpholino (Marlow et al. 2012) and translation blocking 𝛃-catenin morpholino (Leclere et al. 2016) (Gene Tools Inc., USA) were injected into embryos at 750 μM, 250 μM and 1 mM respectively for the experiments. For organizer gene overexpression experiments, plasmids encoding the expression on Wnt1 and Wnt3 (Kraus et al. 2016) driven by ubiquitous promoters were microinjected into zygotes at 20 ng μl−1 each. All microinjections were carried out with 0,125 mg/ml of fluorescent Dextran-Alexa488 or Alexa568.

### Transgenic Lines

For the generation of the mesodermally expressed reporter line 𝛃*-Laminin::eGFP-CAAX*, we cloned a 4.6-Kb fragment upstream of the translation start site of mesodermally expressed gene Beta-Laminin (gene model: NV2.1719) into a PCRII-TOPO vector, driving the expression of eGFP conjugated to a membrane tag and followed by a SV40 Polyadenylation signal. This construct was then injected into wildtype zygotes as described (Renfer et al. 2009) to generate mosaically transgenic F0 animals which were then crossed to generate fully heterozygous F1 animals. These were then intercrossed to generate F2 animals that were then used for the experiments. Refer **Table S4.** for the sequence of Beta-Laminin regulatory region cloned.

### Generation and analysis of single cell RNA sequencing (scRNAseq) libraries

Samples either after drug treatment or post-aggregation were collected in tubes. They were then washed with ice cold Calcium Magnesium Free (CMF) seawater briefly and the samples were allowed to settle. The CMF was then removed and they were resuspended in ACME (ACetic-MEthanol) prepared as described previously (Garcia-Castro et al. 2021) and tubes were left on the rocker for 20 min. The samples were then dissociated using a cut pipette tip every 5 min during the 20 min window and their dissociation status was checked until sufficient. Tubes were then spun at 400 rcf for 4 min at 4℃ and the supernatant was carefully removed without disturbing the loose cell pellet. The pellet was then resuspended with 1xPBS + 0.5% BSA + RNaseOUT (40U/ml) and DMSO was added to be 10% of the final volume. This cell mix was then stored at-80℃ until the day of library preparation. The tubes were then left to thaw at RT and then spun again at 400 rcf for 4 min at 4℃ after which the cell pellet was resuspended in 1xPBS + 0.5% BSA + RNaseOUT (40U/ml) and the viability and concentration were then quantified using ViaStain™AOPI staining solution (Nexcelom) and the Cellometer X2. All processed samples had a viability of at least 85% and raw sequencing data from captured cells have been deposited into the GEO repository *(in submission*)

Sequencing reads were mapped to the *Nematostella vectensis* genome (Zimmermann et al. 2023) using standard parameters of Cellranger v7.1.0 (Zheng et al. 2017). Filtered matrices were then used to proceed with analysis using Seurat v5 (Butler et al. 2018; Stuart et al. 2019). The libraries were filtered for cells with low read and feature counts and analyzed using our scripts (complete details at https://github.com/technau/Gastruloids).

## Acknowledgements

We would like to thank the following funding agencies; Standalone grant of the Austrian Science Fund (P34404), Austrian Science Fund DocFund-Scorpion and the Research Platform-Sincerest of the University of Vienna (369100). We would like to also thank the following people: Lukas Hille for his help with imaging the gastruloids, Alison G Cole for help in generating the single cell RNAseq libraries, as well as Grigory Genikovich for support in generating the 𝛃*-Laminin::eGFP-CAAX* reporter line. We would also like to thank the Core Facility for Cell Imaging and Ultrastructure Research of the University of Vienna, as well as the BioOptics facility of the Research institute of Molecular Pathology (IMP) for providing access to their confocal microscopes. For the purpose of Open Access, the authors have applied a CC BY public copyright license to any Author Accepted Manuscript (AAM) version arising from this submission.

