## Supplementary Information for "Notch coordinates self-organization of germ layers and axial polarity in cnidarian gastruloids"

### Supplementary material

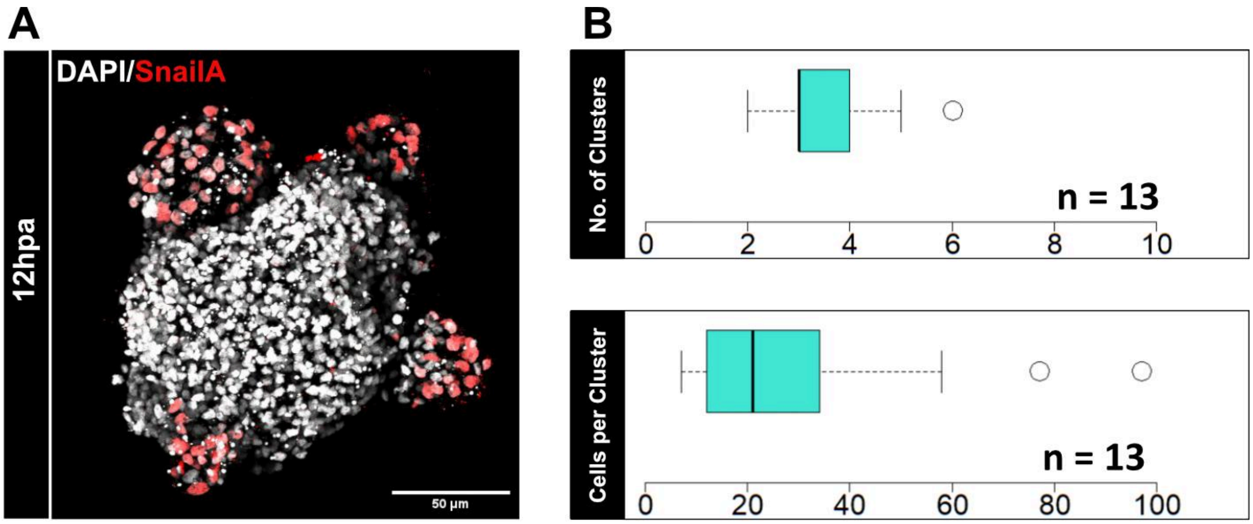


**Figure S2.1. Quantitative analysis of mesodermal cluster metrics just prior to ingression.**

**(A)** Antibody staining for SnailA demarcating peripheral mesodermal clusters at 12hpa. DAPI staining marks the nuclei (Scale bar: 50µm)

**(B)** Cluster metrics quantified from images show on average 3 clusters per gastruloid and 20 cells per cluster per gastruloid.

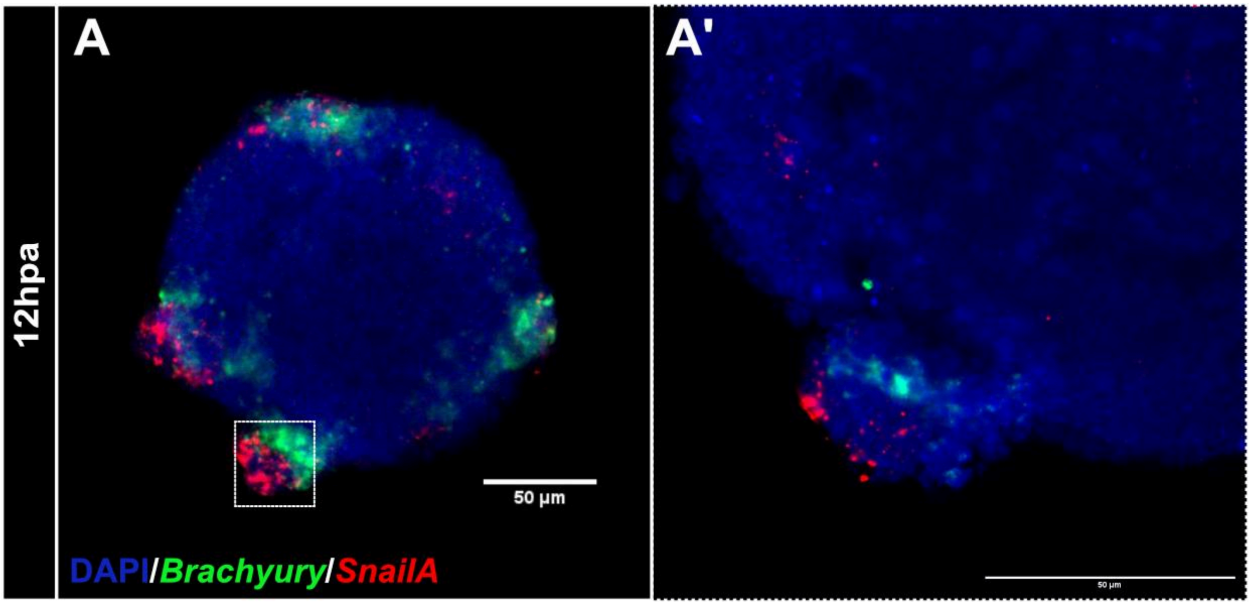


**Figure S2.2. *Brachyury*+ cells form a clear boundary and exhibit no intercalation within the mesoderm.**

**(A-A')** dFISH analysis of *Brachyury* (Endoderm) and *SnailA* (Mesoderm) in 12hpa gastruloids. (A') closeup slice showing clear boundary between both domains (Scale bar: 50µm)

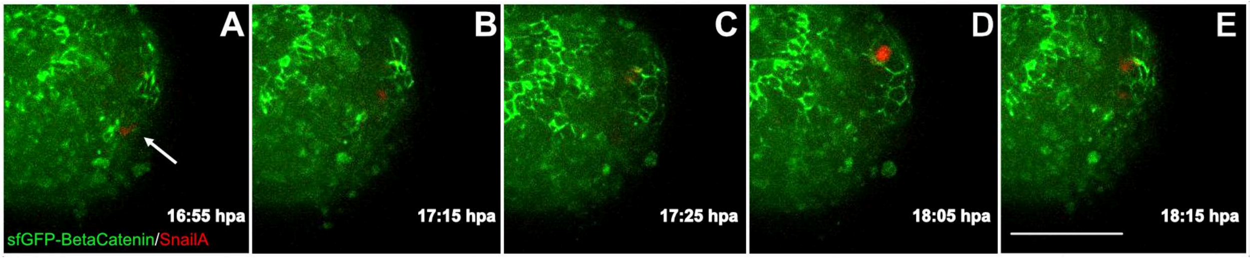


**Figure S2.3. SnailA+ cells ingress at regions of non-epithelialization.**

**(A-E)** Time Lapse images of transgenic SnailA+ cells sorting in gastruloids with cell junctions marked by sfGFP-BetaCatenin. White arrow marks cell ingression at point of non-epithelialization. (Scale bar: 100μm)

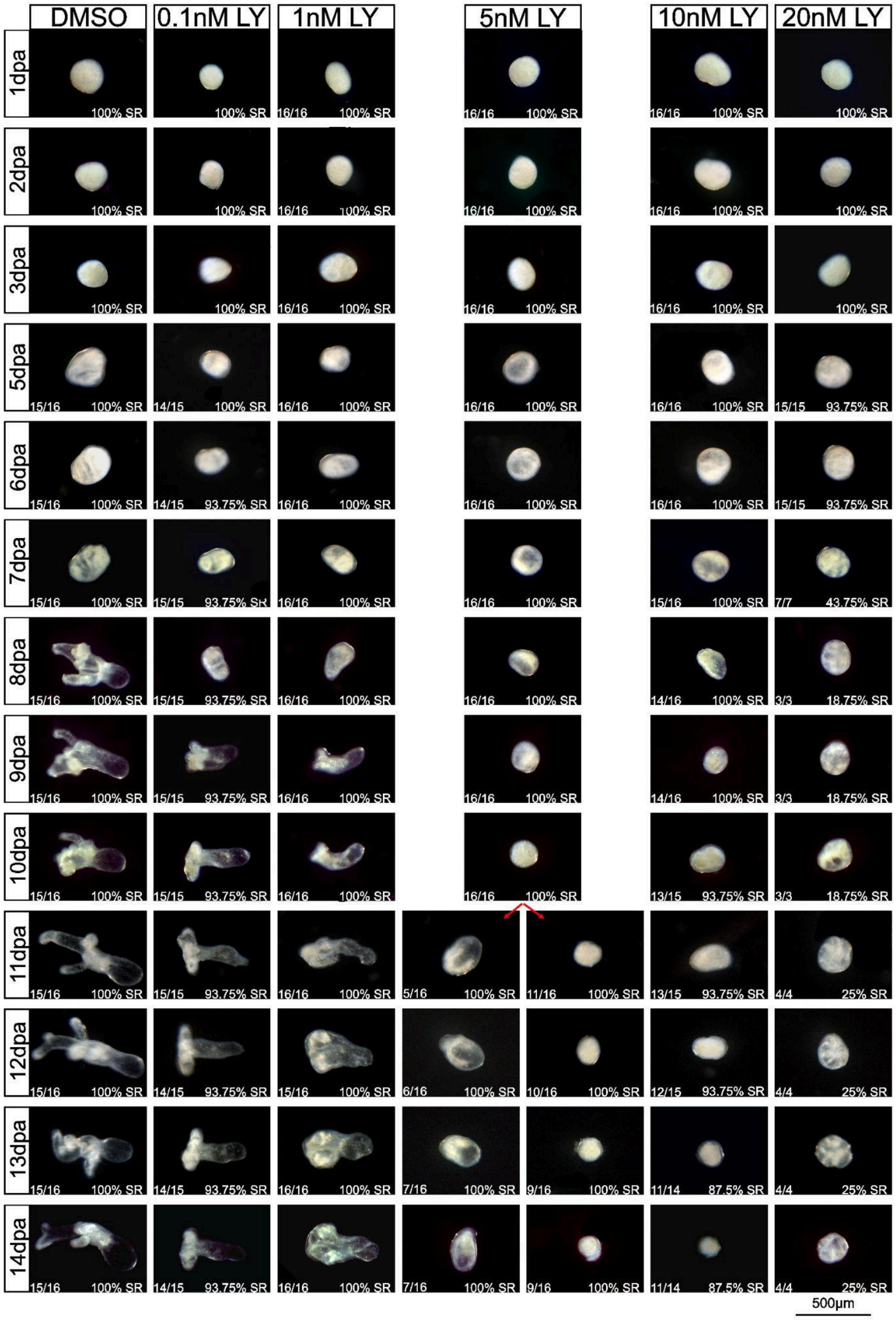


**Figure S3.1. Notch signaling is essential for gastruloid head formation.**

Dose dependent loss of gastruloid head structures (tentacles and pharyngeal tissue) with LY-411575 treatment until 14dpa. Red arrows in the 5nM LY-411575 treatment column represent a split into dual phenotypes. Fraction on the bottom left of each picture represents no. of gastruloids with a particular phenotype out of total. dpa;days post aggregation.SR;Survival rate (Scale bar: 500µm)

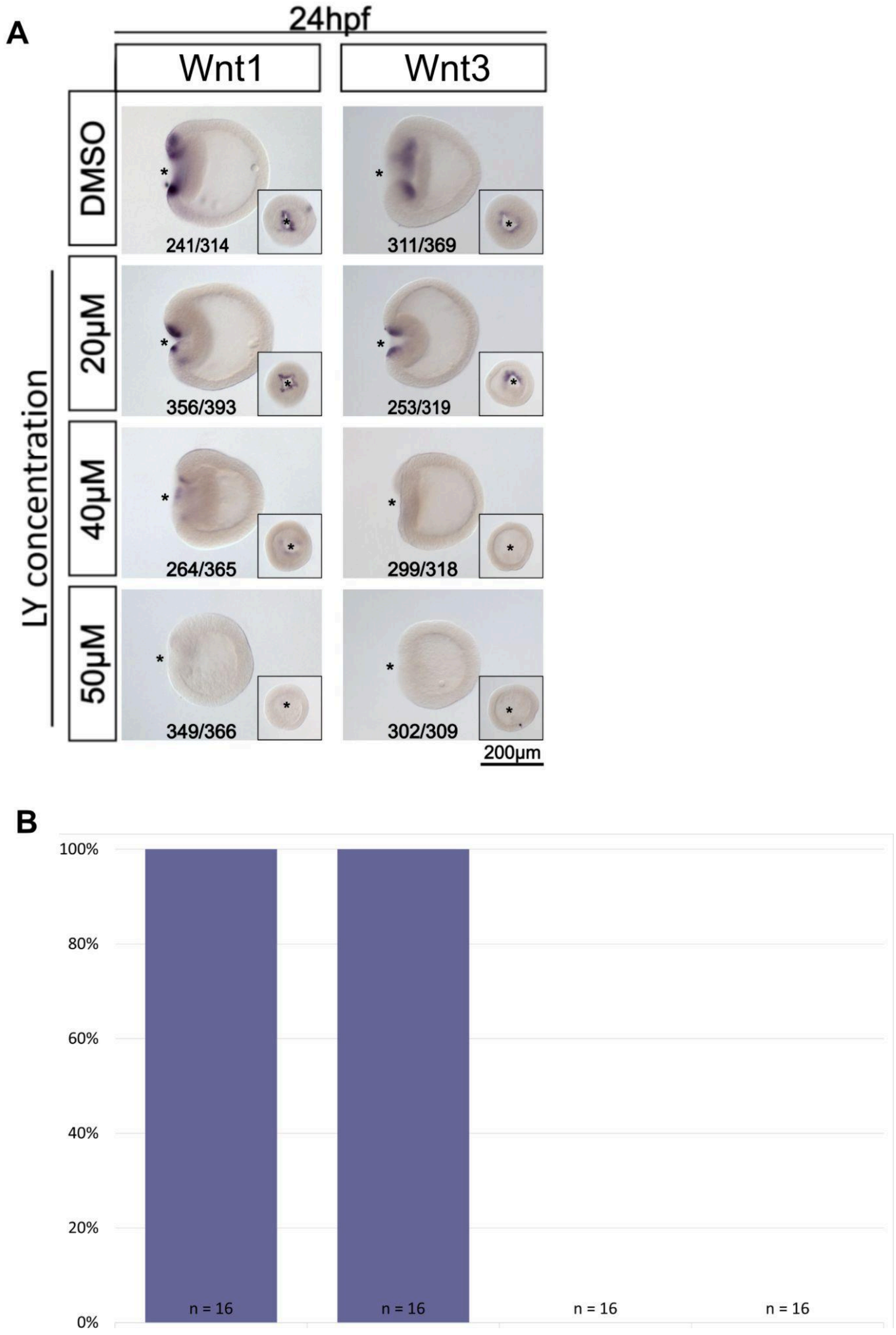


**Figure S3.2. Pre-existing endoderm is essential for axis reestablishment.**

**(A)** Gradual ablation of axial organizers Wnt1 and Wnt3 at 24hpf by different concentrations of Notch signaling inhibitor LY-411575. Ablation was visualized by *in situ* hybridization. (Scale bar: 200μm)

**(B)** Effect of different levels of pre-existing organizers at 24hpf on gastruloid head formation at 10dpa.

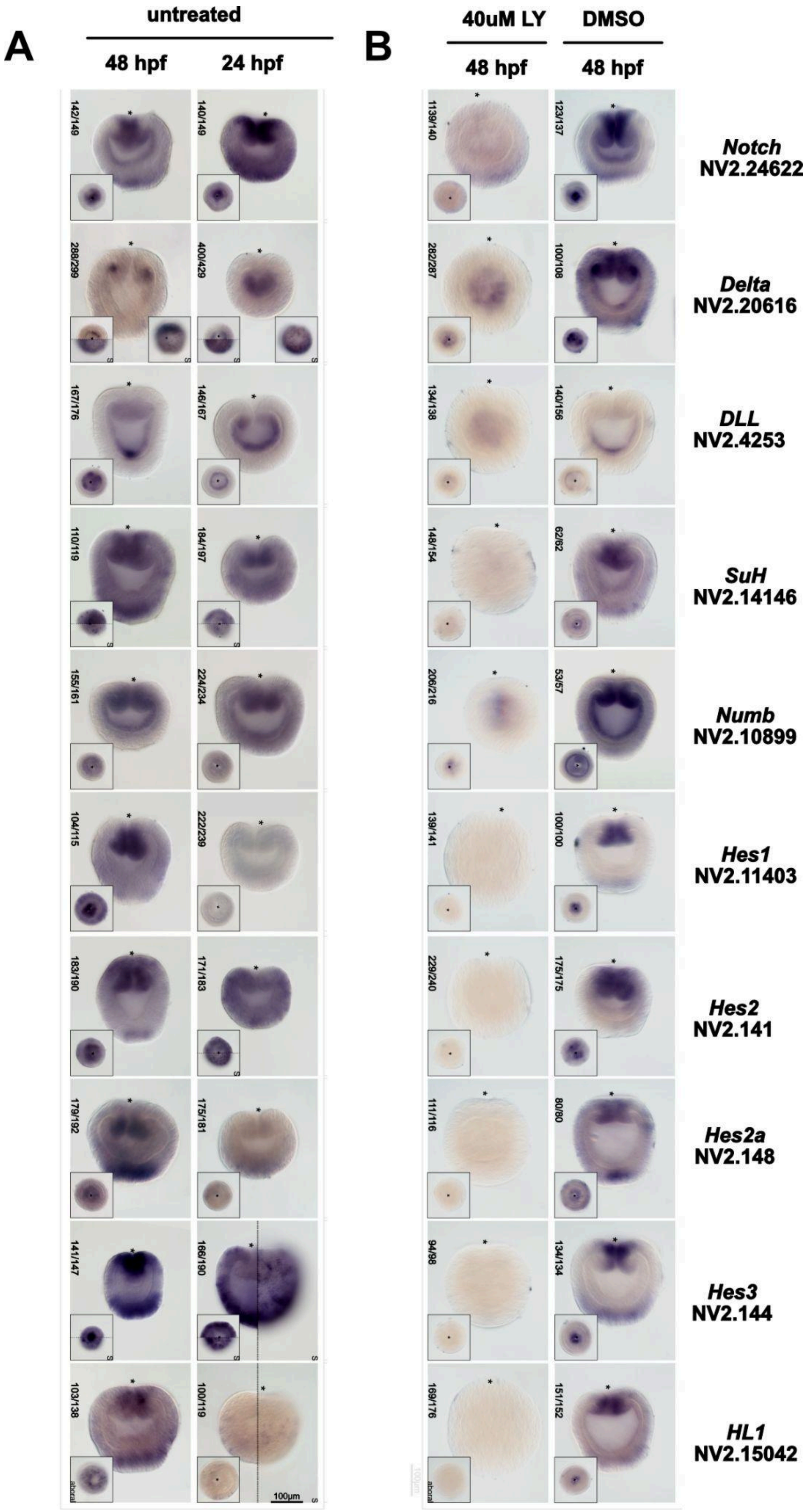
A
untreated
48 hpf
24 hpf
B
40uM LY
DMSO
48 hpf
48 hpf
Notch
NV2.24622
Delta
NV2.20616
DLL
NV2.4253
SuH
NV2.14146
Numb
NV2.10899
Hes1
NV2.11403
Hes2
NV2.141
Hes2a
NV2.148
Hes3
NV2.144
HL1
NV2.15042
100µm

**Figure S3.3. Characterization of canonical Notch signaling components.**

**(A)** Characterization of annotated Notch signaling components by *in situ* hybridization in 24 and 48 hpf wild type embryos. S represents the surface view of the embryo.

**(B)** *In situ* hybridization expression analysis of Notch signaling components in 48hpf embryos treated with DMSO and LY-411575 from 24hpf to 48hpf. Asterisk represents the oral pole (scale bar: 100μm)

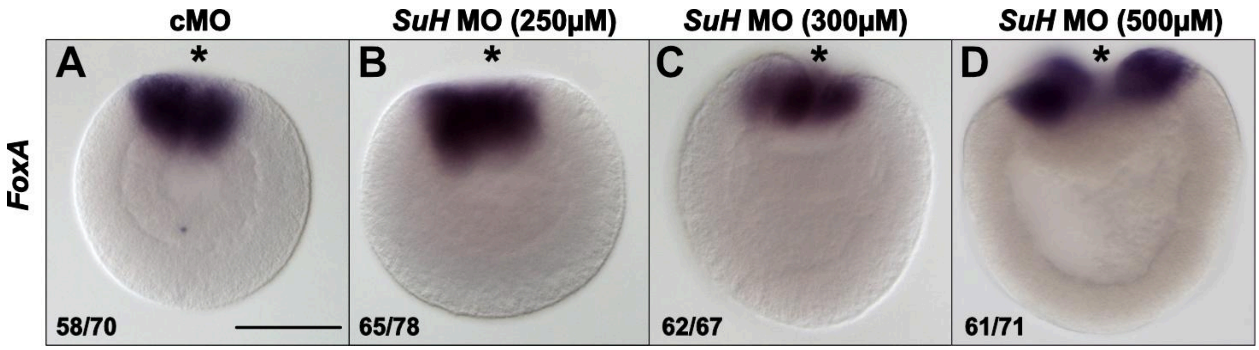


**Figure S3.4. Low dose *SuH* MO injected embryos display no gastrulation delay.**

**(A-B)** Low concentration of *SuH* MO (B) injected embryos exhibit no phenotype and is comparable to the control morpholino (A).

**(C-D)** Higher concentrations of *SuH* MO causes delay in gastrulation.

Asterisks denote the oral pole. (Scale bar: 100μm)

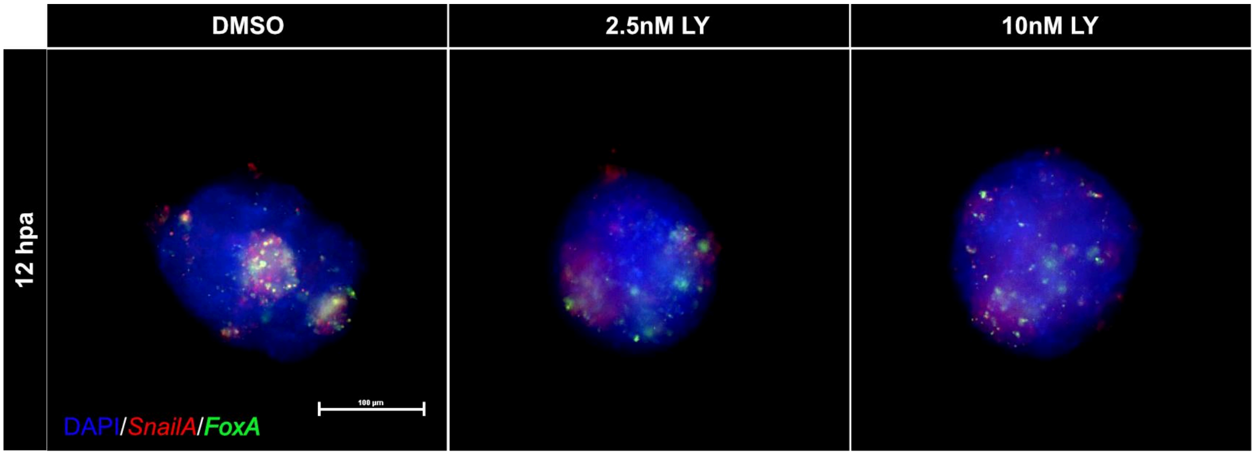


**Figure S3.5. Notch signaling maintains endo/mesodermal boundary in gastruloids.**

dFISH analysis of endoderm (*FoxA*+) and mesoderm (*SnailA*+) boundary formation in notch signaling inhibited gastruloids at 12hpa (Scale bar: 100μm).

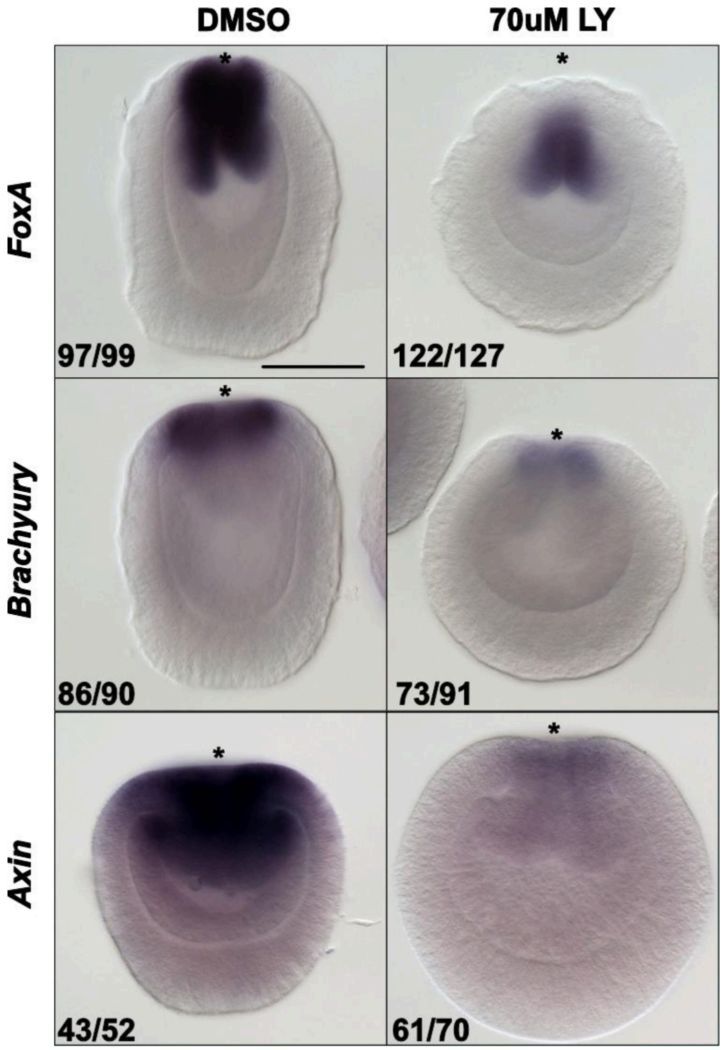


**Figure S4.1. Notch signaling maintains endoderm identity post-gastrulation.**
*In situ* hybridization of endodermal genes FoxA/Brachyury, and BetaCatenin target *Axin* in 48hpf embryos treated with DMSO and LY-411575 from 24hpf to 48hpf. Asterisks denote the oral pole. (Scale bar: 100μm)

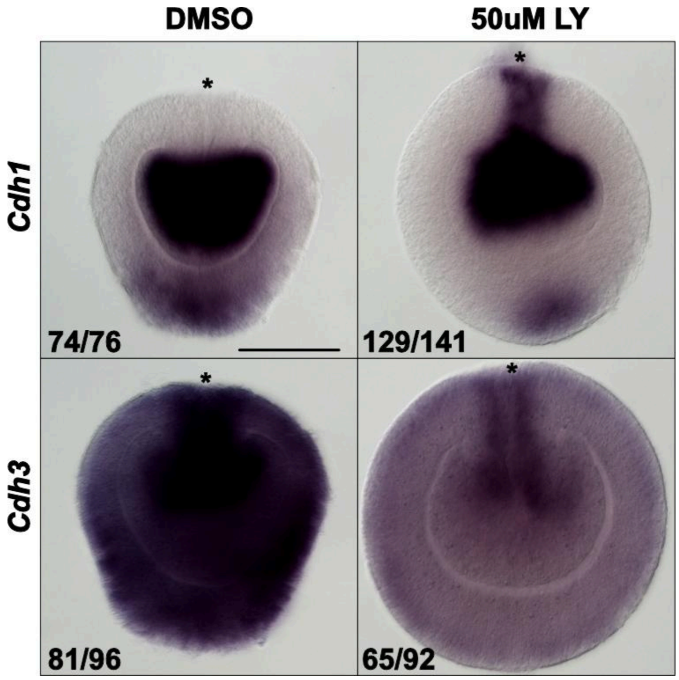


**Figure S4.2. Notch signaling maintains Cadherin Switch.**

*In situ* hybridization of *Cadherin1* and *Cadherin3* in 48hpf embryos treated with DMSO and LY-411575 from 24hpf to 48hpf. Asterisks denote the oral pole. (Scale bar: 100μm)

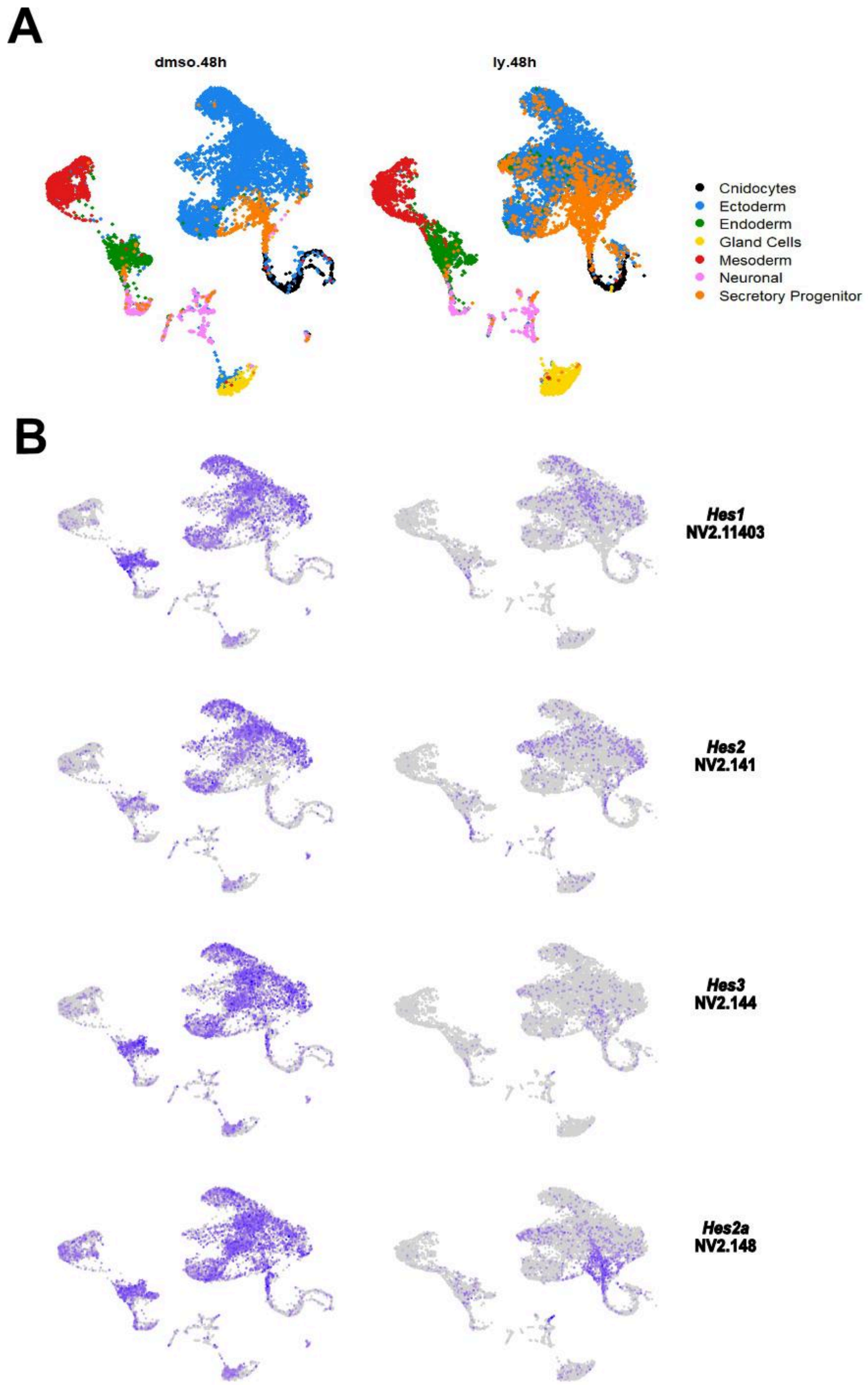


**Figure S4.3. Knockdown of Notch signaling targets in LY-411575 treated dataset.**

**(A)** UMAP dimensional reduction showing annotated clusters in libraries capturing response of embryos to Notch signaling inhibition, treated from 24 to 48hpf.

**(B)** Feature plot depicting knocked down expression of previously reported putative targets of Notch signaling in LY-411575 treated embryos compared to DMSO treated control embryos.

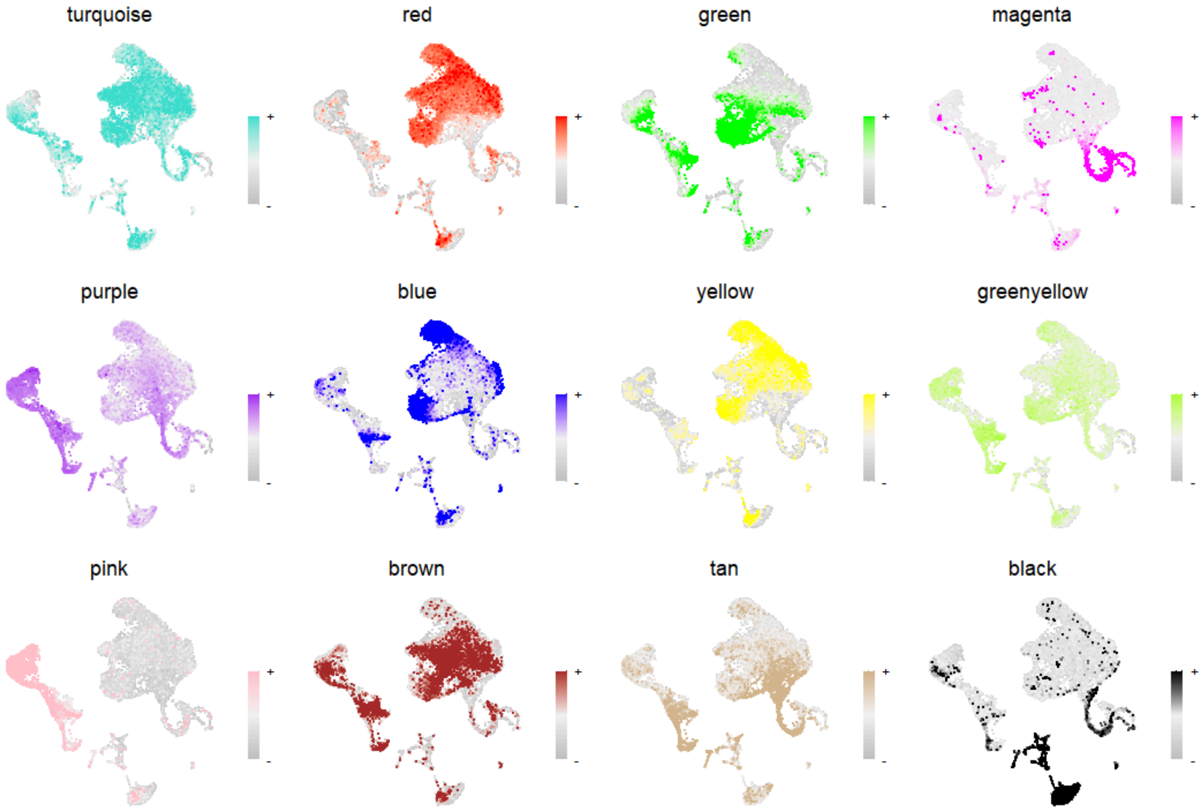


**Figure S4.4. WGCNA gene module analysis.**

Feature plot showing eigen gene expression of identified gene modules

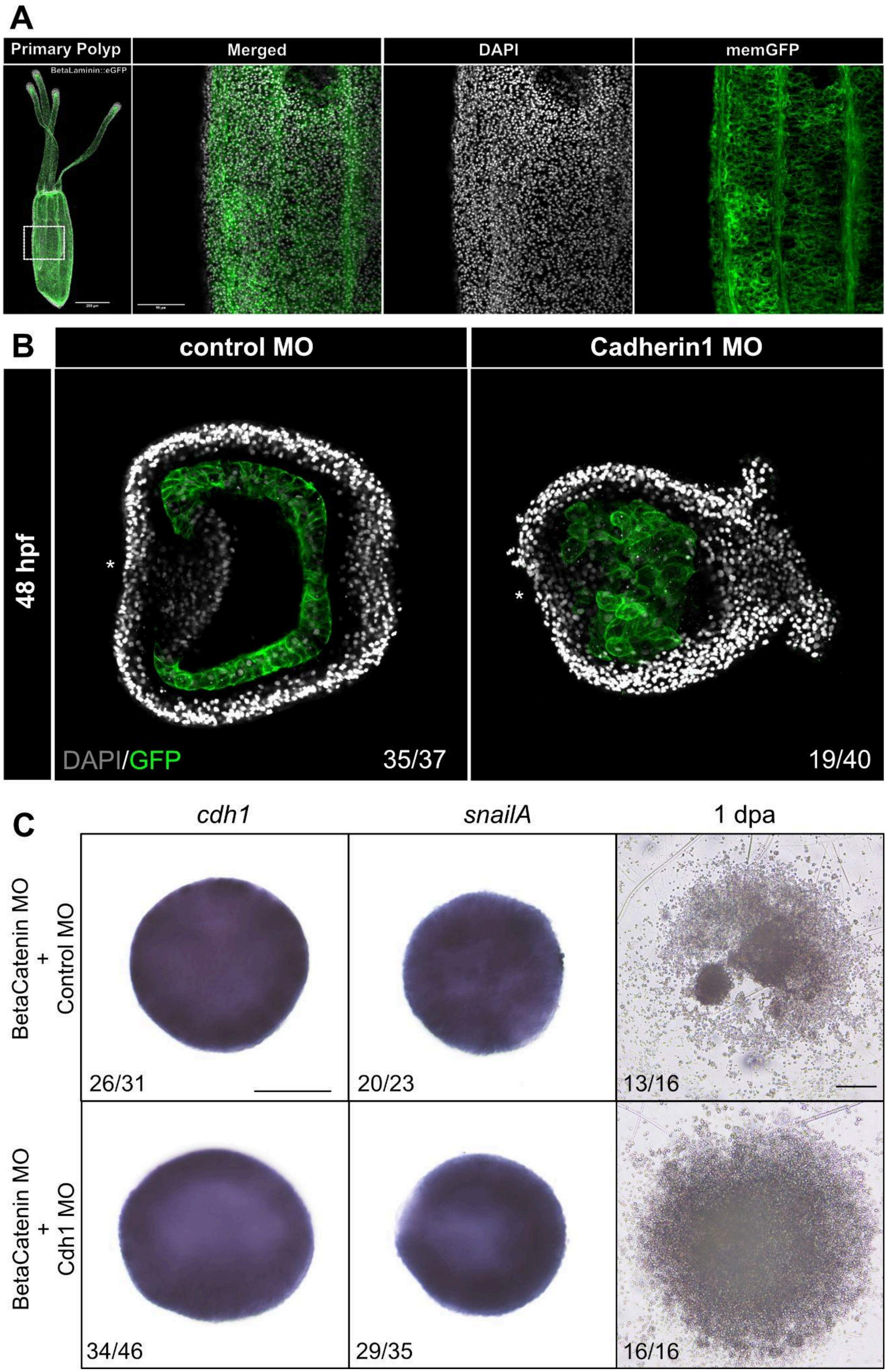
A
Primary Polyp
BetaLaminin::eGFP
Merged
DAPI
memGFP
B
control MO
Cadherin1 MO
48 hpf
DAPI/GFP
35/37
19/40
C
cdh1
snailA
1 dpa
BetaCatenin MO
+
Control MO
26/31
20/23
13/16
BetaCatenin MO
+
Cdh1 MO
34/46
29/35
16/16

**Figure S4.5. Mesodermal adhesion is mediated by Cadherin1**

**(A)** F1 generation primary polyp of β-*Laminin::eGFP-CAAX* line displaying membrane fluorescence in mesoderm.

**(B)** Anti-GFP antibody staining of β-*Laminin::eGFP-CAAX* line embryos post knockdown by control and cadherin1 morpholino at 48hpf.

**(C)** First 2 columns denote expression of mesodermal markers *cadherin1* and *snailA* in embryos co-injected with β-catenin MO/control MO (row1), and β-catenin MO/cadherin1 MO (row2). Final column represents aggregates made with the respective injected embryos 1dpa.

| Common Name | Gene ID | Primers | Sequences |
|---|---|---|---|
| Notch | NV2.24622 | F | AGCTGAGAAGATTTGGTTCATTG |
| | | R | TTGACCAGTCTGATAATAACTCCA |
| Delta | NV2.20616 | F | CAATCGGTACACCTGCTC |
| | | R | CAAGTTCCATCACCAAATGC |
| DLL | NV2.4253 | F | GTCTTTTGGAGTCGCTGTGG |
| | | R | GCTCCCGTTAGAATCACACG |
| SuH | NV2.14146 | F | AAGATTTCACTTCGACAGCT |
| | | R | GGACCCATACCCTCATAGAA |
| Numb | NV2.10899 | F | TCTGACTGCATTCGTGTCG |
| | | R | TGCTAGAGAGGCATTTGACTG |
| Hes1 (HEY) | NV2.11403 | F | CGACGAAAGCCAACCGGA |
| | | R | CAAATTCTGCAATCACCACG |
| Hes2 | NV2.141 | F | CGACGAAAGCCAACCGGA |
| | | R | CAAATTCTGCAATCACCACGG |
| Hes2a | NV2.148 | F | AACTAACTACAACGCTAGCG |
| | | R | TTCAAATTGTCCTCCCCATT |
| Hes3 | NV2.144 | F | TCAACACAAACTCTCAACA |
| | | R | TCGAGGTATTAACTTTTATTGCT |
| HL1 | NV2.15042 | F | CCAAGCTGGAAAAAGCGGAC |
| | | R | GTTCTCTGCTTGGGAAGCCT |
| FoxA | NV2.11441 | F | ATGATGGAGCACACGGG |
| | | R | CGAAAGATTTGATATACCACAACTC |
| Brachyury | NV2.10624 | F | ATGCACTCGGACGAGAAGAAAC |
| | | R | TTAAGCTTGCGGTATGGTGTTCC |
| Axin | NV2.20395 | | nucleotides 1–1123 of Genbank JQ959548 |
| Cdh1 | NV2.4715 | | Pukhlyova et al. 2018 |
| Cdh3 | NV2.4928 | | Pukhlyova et al. 2018 |
| SnailA | NV2.472 | F | ATGCCCCGCTCGTTTCTAG |
| | | R | CTATCCTTGTGACGGGCA |
| Wnt1 | NV2.12225 | F | ATGCAACGATTCAGCGCAGCGATC |
| | | R | TTATAAGCAGTTACTAATGATCC |
| Wnt3 | NV2.12463 | F | ATGAGAGTAATTATCACTGCGATTG |
| | | R | TTATTTACAAGTGTAGATGTTAACCACTT |

**Table S1. List of primers for in situ probe synthesis**

**FoxA (5‘ to 3‘)**

AGGAGGTAATCTGCACAAGTCGCTAGCGGAGACACAGTGAACACCAACACCAGCAGCGCGCAACGTCAACCGGCAAACTGGATCCTCAGTACA
GAAACCTCGAGATAGAGATGATGGAGCACACGGGGGTGCCTCCAGCGGCCATGCAAGACCCATCACAAAACCCGCACGAGCTCAAGAAATCCA
AGGACAAGGAGAAAGCGTATCGCCGGAGCTACACGCACGCCAAGCCGCCATATTCATATATCTCACTCATCACGATGGCCATTCAACAGAGCCCA
AACAAGATGCTCACACTGAGCGAGATCTACCAATTCATCATGGACTTGTTTCCCTACTACAGGCAAAACCAACAGCGCTGGCAGAACTCTATCCG
GCACAGTTTATCATTCAATGATTGCTTCGTGAAGGTGCCGCGCTCTCCTGACCGCCCCGGGAAAGGCAGTTACTGGACTCTCCACCCGGACTGC
GGTAATATGTTCGAGAACGGGTGCTACCTTCGCAGGCAGAAGCGCTTCAAAGCCGAGAAAAAACCGGACCTGAGTCACCTTAGCAAGGTGAGC
AGTATGACACACAACCCGGTCACAGTACAAAGCATGGCGAAGAGCATGGCGGCACAACCTCGCTCAATGGGAACCCCTAGCTTCCTTGCACCG
TCTCCGTACGGGACGGCTATGGGCATGGGACACGTTGGGAGCATGACCGCCATGGGTATGGCAGGTATGCCCATGAATAAGTCGTTTAATCACC
CATTCGCTATCAAGAATATCATCGCGCAAGATCACGAAGCTGAGCTTCGAGGCTACGACCCCATGCACTTCAGTCCTTATCATCCATCACTCCAAT
CCATGGGTTCCCTAGGACTCCCTAAATCCGCCTACGAATCGCAACCTATCACGACGGATACGAGTCCGTACTACCAGGGTTGCGTCTTCACGCC
GTCGAGTTGTGGTATATCAAATCTTTCGTGAACTTAGCGCAGCACTTAGAACACTAGAAGTAGAAATTATTCTAAGAAAAGCTGAATATTATGATAAA
CCTGTATATATAAACATTGAGGCACTGAAAATTGAGCTATCCGACGAATATCTCGTCTATGTACAATGTGCTTGAGCTTTCGTATGATTATATTGATCA
TTTTTTTACTACAAGCACAACTAGATTTCGTAAAGAGCTAGAAAATTTAATAATTTTATGAACAATCAAAAATGTAAAATAGAAGATGAAAATGTTAGG
CTAGGAAAAACAAAAATGATTAGAGTATTTTGAAGAAAGCAAATCTTCCGAAATCAAAATATTTTGAAATTGAATTAACAGACACAAGTGATTATTAG
CTAAGAAAATGTTTGTAAAGATTCCTATATACACATATATATTGTTTCGCAAACTTGTTCTGTTGAAGCTGACCATTAGATGAAAGAGCGTGCAGTTA
ATATTTGGCTTTTCGACAGTCGCTTTCTGTATTCGGGGCATAACATTGACTGTTGCTCAGTTTGTTCTCGTGTCAACTCTGTATTTATCATTGCTTCA
CTTTTATTTATACTAGTCAGGCGAGGGAACGGTTTTTATTGTAAATGTACTCTCATGGTCAACGTTTGTGGATACTTGAAGAAAATAAATTTTTAACG
AATC

**Cdh1 (5‘ to 3‘)**

GGCTGTTATCATGATTATTTTTAGTGAAAGTCACCGCGAGAACTAACGCTTTCTGGCTGATGTTGGCATGACCTTTGCAGTAATCTAGACGCCATT
CAAACGATATCCCAGGCTTAATCTACATTTCCGACCAATCCGAAGCCAGCGATCAAGGACTTTATAAATTCTCAAGCGAGTTATTTTTGCGTATCAA
GTTAGGGAGTCTTCGGTTCAACCTTGTAAGGCGCTCGTTCAACGGCCATATGCTATGGGTTTTTAAACTCACGGGTTCCTAATAACTAGCCACAAA
ATGAGTGCTGGCCGGCTGGCGGCTGTGCTAACCACTCCACTATTATTTCTTAGTTTGCTTAAAACTTTTCAACTGGCAAAAGCGCAGGACACTTT
GATCGAAGTGAATTTCGATGAGGGTCGACCGGCTCGTTCTTCTGTCTACTTGTTCGACTCTTCTAGTGGCGATGTATTCTCGCTGTACCAAGCAG
ATCCGACAGTGCCGCTGCTATTTCAAATATCTGAGGTTGGACACGTAACATCGACTCAAGAAATCGAGTACGAAATAGGAAAAACAAATAAATACG
ACTTGACAGTGTTACAAAGACCTCGAGGAGAAACTCTCGGCGGAATTGCCATAACTCTTCGGATTACCATTCTTGATGTGAATAATTTCCATCCTG
TATTCCAACTCTCACAAGGGGAACATTACGAGGGCTTCGTGAAAGAGGGTACTGCAGAGAATACAATTGTGGAAGGGCTAGAACAGTGCCATGC
AACAGATCGAGATACATCGGGGATACGAGGTTATTCCATCATTAGCGGGAACGAGAAAGGCTACTTCAAAGTCGAGACCGTACAGATCGGCAGC
GGGGTAACGAGCCGTAAGTTCTTAGTGTTGAAGACTACGGGTAAACCGATTGTTCGTGATGATAATAATCCTTATATTATGCTGACGGTACAAGTCA
CGGACGGAGGGAACCCCTCGCACTCGGGAACTGCGAACATTCGCGTGAATGTTGAAGACGCGAACGACCAGACGCCGGTGTTCGAGAGCTCA
CAATACCGTGAAACAATAGCCGAGAATACTCCCATTCAGACCTCTGTGCTACGCGTACGGGCTACAGACAAAGACGACGGCACGAACGGCGGGA
TCTACTATTACATGAAGAATCCTGTGAATAGCTATTTCACCATTGATGCTATTACAGGAGTTATTAGGGTCGCTAAAACGCTCGACTATAACGCCAGG
GATAAACACACGCTGTACATTTCCGCACGGGACCGGGGGGACCCTCCTAGGACGAGTGCTGAAGCCACCGTAGAAATTTCACTGAGAGAGAATA
TTCAGGGATGGCCGCTACCTGATAGCGCAGACCCGAAGGAAAATACGAAACCATATTTCCCCCAGTCCAGGTATACATTTAGTATTCGCGAGGATT
TTCCGCCTAAAGGTGCATTACTGGTGATGCGCGCGGCGGACAATGACCCGATTGGACCAAATAAGAGATTACGGTACTCTTTGTCTGGAAATGGT
GTCTCTAAATTTGCGATCGATCCTGAGAGTGGCGTGGTAACACTGACGGATTCTGTGGACTATGAAAACACCCCAAATAACCACGTGTACGATCT
CACAGTCACCGCTACCGATCAGGGCCCAGGGTCTCTTAGCGCCACTACACAATTACTCATCGAAATCCAAGACGTCGACGAGAATAAAAACTCCC
CAAGGTTTGACCCCCAGCAAGCAATCCTGGAAATTAGTGAAAATCTTAAACAGAACTCGCTAGTGACTACAGTAAGCGCTACGGACTCAGACAGT
GTCGGGAACCCGTCGAGCCCTGATGGGAAAGTGGTCTACTCGATTGTAGGGGGAACTGGGCTTGGCGTCTTTCGCGTGGATTCAAATACAGGA
GAGGTGAAAGTCGCTGTACCTTCTCTAGACAGAGAAGGTACATCCCAATACACTCTAGTTGTGAAAGCGAGCGATAACGCTACATTCCCACGATC
GTCGCGCCTTTTTCTCATGATAAATCTATTGGACGAGGATGACAATTTCCCGTACTTCAGCCAGCCTATCTATATAGCCCAAGTCCCGGAGAACCA
GCCGTCTGGGACCTTCGTCACAGTGGTGGTTGGCAGGGATGCGGACGAGGGATTCAACCCAAGCTACACGGTGATCACTCCAGGGGTTCCGT
ACAAAATCGAGCCGTCAACGGGCGTAATCCGGACGTCCAAATCTTTGGACCAATCAGAGTTGAGCACAAGGGAGCTGCAGGTGATTGTTCGCGT
TAGTACTGCCGGGTCCAAAACAGCGGACGGACAAGTAAATATAACAGTAATAAGCAAGGTCAACCGTCCACCCGTGTTCAAGAATACACCATATTC
TGTGAAAGTCCCCGAGGAAATGGGGCCTCTGCCCAACTTATTCTGCATCGCTGCTGTTGACAGCCTTTCGAGGCCGGTCCAGTACACCCTAGCG
CCGGGAGCAGATGGTCTATTCGAGGTCGATAAGGACTCCGGTCGACTGCATACCAAAAACTCCTTCAATTACGAGAACGTCGATCGCTACAATCT
CCGGATCGAGGCTCGCACCAGCTCCGTCCAGGAAGTGGCGTCCGCCTCGCTCACCGTCGAGGTCACGGAAGAAAAAGACCTGCCAAAATTCT
CTTCCGACTCGTACCAGCTAACTGTGGATGAGAGCGCTGCACCGGGGACTACCCTGGCTCCAGGGCTCCTCATCATCGACTCCGACACCTCCA
GTGATCAATTCGACTGTTCTATGGAGGCTATCACCTCTCTCCACACCCTATACAACTTTGAAGTCACCCAACAATCAGGGCGTTGTTTTCTACGCG
TTCAAGCAGGGGGAAAGCTAGACGCTCATTTAGCATCCAAGTACACGTTCAACGTTCGCGCGACTGACAGGAATTTCAGAAACATGTTTGCGAC
AGCCCAGGTGGAGGTCAACGTGATTGACGTGAATGACCATAAGCCGGAGTTCTTGCAGGAGTCGTATTGGCTTTCGGTACCCAGTAGTACCCCC
GCGGGGAGTAGTTTGGTGACGGTTCAGGCCGAGGATATGGATATTGGAACCAATGCGCAGGTTCGGTATGAGCTGCTGAGGCAGGAAAACTCA
GAAAGGTTTATCTTGAATGACAACAACCAGCTGTCCACAGCGTCCACGCTGACTCCGAACGTCCGTTACCAGCTGCTTATACGCGCCTCCGACT
CAGCTACAAGGAACCCCTCCTCAGCCCAGGTCCCCGTCTACGTCTCCGTGTACAGCCCAAGTGAGTCTCCGATCGTCTTCGACAAGTCTTCGTA
CAATCAAAACCTTCCCGAAGATTCTTCTGCTAATACACTTGTATTCACGGCTAAGGCCACGAGATCCGGCAGCAGTAGCGGGATCACGTACGAAC
TCGTCGGTGGTTACAAGCAGATCGGCGAGGCCATGTTTAGTATCAAGCCGGACACTGGCCAGGTCTACCTAATAAAAAAACTCGACTTCGAGAC
CAAGTCATACTTTCCGATTGCTGTCCGAGCCAAGTATTCGGGAGGCGCCATCGAGCTCGCCTCCGAAGTAGTAGCCAAAGTCACAATCGTCGAC
GTCAATGATAACGGCCCGAGGTTTGCATTCCATGAGAGCAGCAAGACTGTAGTGATTGACAGCTTTTCTGCCAAGGACACACAATTGGTCCAGG
CACGAGCGCTTGACGCTGATTCTGGTAGCTTGGGCGAAGTGACGTACGGGATCGACGGTGGCAGAAGTACAAGTAACTTGCCTTTTAATATCAA
CGTAAAGACGGGAATGATTTTCGCCACCAGGGAGATACTGTACACACAAGGGTCATCGTACATAATAATCGTCGTGGCAACAGATGGTGCCACGG
ATGGTAGCCAGAAAATCCAGAAATACACCGTCAATGTCCAGGTACTGGACACTCCCCGTCCGCCCAGTTTCCCCCAAAAGACATATTCCGCACCC
GTGACTGAGACTGCGGGAGTTGGGGACACCGTGACCACCGTCAGGGCAGTGTACAGCAAGCCTAACGCCTTCCTCAAGTACACCTTCGTGAGT
GGGAATGAGGACAACACCTTCTGTGTCAACGGGTTTGGCATCATATCTGTTGCCAAGTCACTGGACCGAGAAAAGGTCGCTGGGTACACGCTG
GGCATGCGGGTGACCTTGGGTCAGCACGTGGACGACACTACGGTGTACGTGAACCTCACTGATATCAATGACGACGCTCCACACTTCACATCCG
CTATCTACAGGCGCTCCATCAAGGAAGGACTTGCTGAAGACACAGAGATACTGCCTCCTGTGATTGCTGTCGACCATGACTTTGGCAGCAACGG
AAAAATTTTGTACTCAATCCTTTCCGGGGTCCATCCGGACTGGGACAAGTATTTCAATATTGATTCTGCGACTGGTAAAATCACGACTAAGATGACA
CTGGACTATGAAACGCACAAGTCGCACACGCTGTTTATACGAGCCGAGGATAATGGCAGTCCCAAGAGGCTCAGCGGAATTGCTCAAGTCGACA

TAGACGTGATTGACAGGAATGACAACTCCCCCGTGTTCGCTGCCGCGTTCTACCGCGCAAAGATCTCCCTCGGTGCGGTCAAGGGGACGAGTG
TGCTACAAGTGCATGCTACTGATCTGGACAGCGGACAGAACGGTCAAATAACGTACTCAATCATCCAAGGCAACGAGGAGGAAGCCTTCACCAT
CAACGAACAGGGTGTCATACTCGTGGACAAGTCGCTAACAACGGTTGCTGCGGATAAGTTCTCGCTGAAGGTGGAGGCCCGCGACAAGAACGC
ATCTCCCCGCAGCGGTAGCGTTACTGTTGAGATCAACGTCTACCTTCCCGACGGTCCACCCATGTTCGTGGTTTCCCCGGTTACGGTGTACGTTA
AAGAGGGCGTCCCAGCCAATCATCGCGTCGCTGGTGTCAAGGCTGCCACGTCAGAGGCCCTGACGTACACTCTCTTGTCTGGAAACGAGGCC
GGCATGTTTAGGATCAACCCATCTACCGGTTCCCTGGATGCGACCCGCGAACTCGACTACGAGGAACGCAGGCGCTACGAACTGAAAATCGAAG
CACGTGACACTCGCGACCGCAGCGCAATGGTTGAGCTTATCATCATTGTCGTCAACATCAACGACAACAAGCCCACCTTTGTCGACATCGTCGA
CGGACAAATAGATCGGAAAGCCATCGGGACGTGCTCGCCGACCGACATTGCTGAAGGAGAAGTCATCACTCGGCTCGGCGCGTTCGACCGAG
ATGGCGACGAGATAACTTTTATAGTCCCGGATGATGTAAAAGACCTCTTCAAAATCGACTCCCGTGGTGTCGTTACTGCCAAGAAGCCTATCGAAG
ACTTACCGAAAATTTACCCGTTTAACGTCAAAGCTCTCGATAACGGCGAACCGCCACAGGAAACGGACGTCAAAGTAAGACTTGTTTTTGTTCATT
ACCGCCCGGACCAGAAGCACGTTCGGGTAAATGTCCGGGAAGACACGCCTACCGGGAGTGTTATCGCCACTGTCCGGAGGTACTTCCCTAACG
GCATCGTCTCATTGTTGCTGCCAGAAAAAGCGAATTTCAGCGTAAGGGCCAATGGTGACGTCATCTTGTTAACACCGCTGGACTACGAGAGCCA
GCAGTTTCACCGGATGACCGTCCGAGAGGAGCTTGGTAACCAGACAAATGATGTTGATGTGGAGGTTGTTGTTTTAGATGTGAACGACAACAGG
CCGATCTGGATGGAGCGAGAGAGACTGGCGCGAGTAAATACTAACAGCCGCGCCGGCGCACAGGTATACCAACTCGAGGCAAGAGACGAGGAT
GACGGTTCCAGTGGCCTTGTGGGATACCAGTTGAAATCTCCCTCGAATGATGCGAAACGCAAGCGCCGCTCACCTCCACAACATCCTTTCACCA
TTAACCCCAAGACGCGCCAAATGGAGGTGGCGGGCTCACTGAAAGATCAACGCTACGATTTGGACGTTTTCGCATTTGATTACGGGATTCCGCG
TTTAACCAGTGACACTATTACGCTCAACGTGGATGGCTCTGCGGCAGGGCAACTCCCGCCCAGGTTCTCAAAGGTCAGCTATCACTTCATGGTAT
CCGAAGACGCGAAGTATCTCTCGCTTGTTGGGATAATACTTGCTCGCAGCATCTCTGGAGCCCGATTGGACTACAAGATCGTCAGTGGTAATGTC
GGAGACAAGTTTATCGCCATGGGTGATGGACGAATCTTGCTGAATTCCTTGCTCGACTTTGAGCGCGACCAGACGCAGTATAATCTTAAAGTGAA
AGCGACCGAGCAAATACCTGATGGTTTGGACTCTACAGTAGACGTCAAAATCGATGTCATCAACGCCAATGATCACTTCCCGTACTTTGATCAGCA
ACTTTACTCGGTCCAAATACCGGAGAGTACTGCAGTAGGAGTCCTCGTGCAAGAGGTGACCGCTAAAGACTGCGACTGCCTCTCTAGCTGTACC
TGTTCTCCAGGTTTCCTTACCTATTCCATTGAACCCAGTAAAGAGCAAGGAGGAAAGTTCTACATTGATCCGGCAACGGGTAAGATTTCAGTCAGC
GTGGCGTTAGACTACGAGGATCAACGATACCACTTACTAAAAGTCTACGCAACGGATAAGGGAAAGAAGTCCTTCCAAGGACTGTGCTTTGTCAA
TGTTACCCTGACTAATGTAAATGACAACCGCCCTACCTTTCTAAAGAGCGCCTATGAGTTCCGCGTAGCCGAGGGCGCCGCAACCGGCGAATCC
CTCGCGATCGTGGTCGCAGTGGACGCAGATGGAGACGCGGTGACCTATTCAAAGGCAGGTGGTGCTTTACAATTCTCTGTGGATCAATCAACCG
GCGTGATCACACTGAACAGTGCTCTGGACCCGAACAAGAACCAGTATACCTTGCAGGTGATGGCCAAAGACTCTGGAGGACTCACAAGTACCGT
ATCCGTCACTTTCAACGTCGCAGATGCTAACAACAACGTACCACAATTCACCAACTGCGGGACAGTCAGCATCCGCGAGAATGACCCGCGGGAC
ACTAAAATCACGCAACTGACGGCTACGGACGCGGACAGAGGGCAGAACGGACAGATAACTTACTCCATTGAGGACGCCAGCTCACAGACTCTC
TTCTCCATCGAGCCCCGTACTGGTGTTGTCCGCAGCCTCACCAGCCTCGACAGAGAAAACAAGGACAGTTATAATGCAATTATCAAGGCAGAGG
ACGGGTCTTCGAAGCAAGATGAATCTGAGAGATTACTTTGGTATTGCTACTTGACCATTAATGTAGAAGACGTGAACGATAACAGGCCGTATTTCC
TCGCCGCTAAGTACTTCGGGAGCGTGTTTTCCAGTGCGCCTAACGGGTCCAATATTCTTACGGTCCAAGCTACTGATGCAGATAGTGGCTCTAAC
GCTAAGATCAAGTACGCATTATTAGACTCTGCTGGGGGGCTCTTCCGGTTGGATTCGTCGGGGATACTCAGAACCAATACTAATCCCGCCCGGCT
ACAATTGGAAACTGGGAAGAAGCTACTTTTGGAGGTGTCTGCTAAGGATGTTGAGTCTATCGCTGGGACGCAACCCGGTAAACCCACAAAGTAC
ACCACCCAGATCGAGATACTCGTGTCGAATGAGGAACCACCGAAATTTTCTCAACAGGTTTATACAGCGAGTATCAACGAAAATATGGAGACTGG
ATCTACGGTCACACGCATCACCGCCACTTCTAGCACGGGCGCAGAGATCAGCTACGAGAACGTTGACACGAATCCTCGCGCGAAAATTCTGTTC
CGCGTCCAGCCGGACGGGTATATCATCACGGGTGATCGGCCCGACTACGAGCGGGGTACCACCTACAACATGCAATTCGCCGCCAAGGACAAA
AAAACGTTGCTTTACTCCACCGTTAAGGTTGTCATCAACATCATAGACGTAAACGACGTCTCTCCAGCCTTCCTCCTCGCGATCAACACACGCAA
CGCGCGCGTCCTAGAGAACAAACCCGCGCCCACGAAGGTTATTAGCATGAAGGCCATCGATGACGATGGCAGTGAACCGCATCGACGCGTGAC
ATACGAGATGAAGGATAACCCCAATTTCCAGATCGACGCGAGCACGGGCATGATCACTACGAAGACTACTTTGGACCGCGAGGTGACGCCCAAG
TATGACGTCGAGGTGACGGCGAAGGACGGGGTCAACAAAGAGTCGGCCATTCTCTACATTACGGTTGTAGATCAGAATGATCAGCCTCCCGTGT
TTGCGCCCAAGAGCTACGCTATCAGTGTTCCTGAGGATTCACCCATAGGTACTTCTGTGTTAGATATCTACGCTACTGATGCAGACGTTGGCGAAA
ACGCAAAAATTACCTACTTCATCTCAAAGGGTGACCCCGAGGGCAAGTTCTCCATCGTCACTTCCCCCGTCAAGGGTGAGCTAGTGGTGAATGG
TAAACTGGACTTCGAGACAAAAAGCAGTTACACGCTTGAGGTGACAGCTACCGATGGCAAGTTTAGCGACACAGCTGTGGTTACTGTAACGATC
CAAGATGTGAACGACCTCCCACCCGTGTTTTCCAGTCCTCTGTACGAGTCTCGTATCCAGGAGAACACAGGGCCTGGTGCGGGCGTTGTCATG
GTAACAGCATCGGACATTGACAGCCCCACAATATCCTTCTCACTGGATGACCGCGGAAAAGACTACTTCCAGATCACGCCCATTCGTGCGAGTG
GCCCAGGCAACGTGTGGGTGGGTGACATTAGGACCGGAAGCAAGCAACTTGACCGAGAGGAGTCGCCTGTCAAGGTGTTCACCGTCATCGCC
AACGACGGCAAGCACACCGCACAGGCAGAAATCAGGGTTAACCTAACGGACGTCAACGATAATGCTCCAAGGTTCCCAGCGTCCCCGTACATTG
GATACGTAGAGGAGAACAAACCGTCCGGGACAAGCGTCATGTACATACAGGCTGTCGATGATGACGACCCATTGGCAGGCGGAAATGCCAAGCT
CTCCTACGAACTAACGGATTCCGCCGGAGACAAATTCAGCATCGACCCTCTGTCTGGACTTATCAAGACAAAGGTGACATTCGACCGGGAGCAG
ACTCCCAATAAGTTCAAGGTACGCGTCAAAGCGACCGACGCTGGAAACCCGAGACTATCTGCAAGCGTGGATGGTATTATCCATGTTTCTGACGC
CAATGACCACAAGCCGAAGTTTACGGAGAAATTTTACCGTGGTAGTGTTGCGGAAAATGCGCCACCGGGATACTCGGTACTCCGAGTAACAGCA
ACCGATGAGGATGTTGGCCCCAACGCGGAGTTTGAGTTTGTGGTGGTCCAGGGCAACGACCCACACGCATTTTACATTGACCCGTTTAACGGTA
CTGTACTGGTGTCGGGAATTCTTGATTATGAGAAAAAGAAAGAGTACACCATAACGCTAACGGTGGCAGACCGCGGAATGCCTCCCTTACAAGGC
GACGAGACCGCTTATGTTGTAATTGAAATTCTGGACGCAAACGACAACGCGCCGGAATTTATACCAAAGATATATAACGCATCGGTGCTGGAGGAT
GTCGGAGCTCGGCAGCCTGTGCTTACTGTGACTGCAGTTGACAGGGACAGCGGTCCTAACGGGAACTTCACATTCGCTATCGACCCCCGAAGT
GATCCCGACGATGCGTTCACAATAGAACCTAATCCAAATAATGCAAGTATCGGAATTATCCGTACGCGCGTACCTCTGGACCAAGAAAAGACTCCA
TCGTTCCACCTTAAGGTTACAGCTACGGACGCAGGTGGACTCCAGGGTGAGGGGGAGGTGAGGATTAATGTAATTGACGTCAATGACAATGGCC
CTTGGTTTGTTCCCCCTTTCTTCGTTGGCCAAATCAAGGAAGGGGTGAGTGCGCGTCAGTTTGTCACCAAGCTAAAGGCCTACGATCCCGACGC
ATTTGTCAAGGACCAAGTCATCACCTTTTCTATTTACAACGGAACAGTGGGCGAAAACTTTAAACTAGACCCCGTCTCTGTGACTAATGAAAGCGT
AGACTTGCATTCTTACGGTGTCTTCGACCGCGAGGCTGCTCCAGTGTGGAAGATCGGCATCGAGGCTGTGGATAACGTTGGGCCGAAACCTCA
GAAGAACTTTACATACGTCTACGTTGACGTTCTAGACGTTAACGACAACGCACCAAAGGACGGCAGCTTGCTTATCATTGTCAACGCCTATGACG
GGAATTTCACCGGAGGTGTCATCGGGAAACCATACTACCAGGATGATGATTTTGACGGCGACGAAAACACCTATGAGCTCAACTCTCAATCGCCG
GGTAGCTACTTTAGAGTAAACGAAGGCAACGGTGACATTACGGCCGCCCCAATGATACCCATGGGGGAGTACAACCTCAAGATTCGTGTTACCGA
AAAGAAAGACAGCCCAAGTACTGTAACGTCATCTGTCCGAGTCCTAGTGCGGCGAATAGACAAAGAAGCTGTCGACAATGGAGTAGCAGTTGAG
TTTACGGACATGAGAAAGGTTGGGTACTTCGTAGGGGATTACTATAAGGGCTTTGAGGACGTGCTTGCTAGCACTCTGGGTGTACCTACTGGAGA
TATCAAGATATTTAGTGTGCAGAAGGCTCACGATAATGGACTTGCTGTAGTTGTGTTCTTTACTGTTGCGGCGAAAGATAGTTACATGCCTCATTGG
GACGTTGTATCCAAACTTGTTGACGCCAAGAAACCGCTAGAAAGCCTCGGGTTAAAGGTCAGTCGACTAGGAATGGATGAATGCTCCAAAGGCA
ATGTCGGTCAAAGTGTTGGCGTAGCCAAGAACATCCTTGTACGGTCATCCAATTTCTCCGTGGCCAGCGGTGATTACGGAAAAGTTCCCGCCCC
AGCATCCTCTCTGACTATCGTTTCCATAGACATCTTACCCAAATGTCTCTACGAGGCCGTATTCCCACCTGAAAAGCGCTGTAAGCCGCATAACCC
GTGTCTCCATGGTGGCAAGTGTTACGAGACGGTGCCTGATTGCCCAGGCTTTGTCTGTAAATGTCCCACAGGGTACCATGGTCCCTTATGTGAG
ATGACAACACGCACATTCTACGGCAACTCGTACATTTGGCTTCCAAAACTCATGACATACTCCTTAAGTGACCTCGAGTTCGAGTTCATGACGAAA
ACGGCGGACGGTTTGTTAGTCTATCAGGGACCTGAGCGTGAAGGTGCAAATAACGGCCTGAAGGACTTTATTGCCGTGGTGCTGCGAGGAGGT
CGCGTAGAGCTCTTTGTTTCACTGGGACTGGATCCTGTGACCGTTAAGATGGATAAGGGACCTAGGCTTGATGATGGAGAGTGGCATACTGTGC
AGGTCCTCAGAAATATGAAGGACATCGAAATCATAATCGACCGCTGCTCAACTGCTTTGCTTGAACACAAACCAGATGGCACGGTGGTGGAGAAC
CGCAAATCGTGTCACGTTTACGGGCGCATGCTGGGAAGATCTGTCTTTCTCGATGGCTTCGGTCCACTGCAGATTGGAGGCGTCTCAAACCCAA
ATATGGATTTCCCCGACATACCATATACGGGATTCAAAGGTTGTGTTCGTAATATCAAAGATAATCATAACCTGTACGATCTTAAGAACCCACTTAAA
GTCGTGAATGCCCCAGAAGGTTGCCAGCTCGCTTCTGCTTGTCCTGAATGTAAGAACGATGGTTACTGCGAGCCGCTCATGGCTAGGGACAGCA
TTTGCGTGTGCAACCCTGGATACAGCGGGAAACACTGCGACGGACGCGGTAAAGCGAGTTACTACCTGGCCAGTAGCTTCACTGAATATCTGGT
TGCTGCCCGTCGTCGTCGTCGTGAGGTTGTTCCACCACCCACTGAGATCTTCAATAGGTTCTACACCACCTTGGCGTTACAGGTTAAGCTGGAT

| |
|---|
| GAGGACGCTACGAATGTTGTGGTATTCCTTGCCTCCAATCGCATGGGAACAGAGTTCCAGAGAGTGGACGTTAAGGACAGTAAAATCCGTTACGT<br>GCTTCGCTTGGGCGCACGCATGCTTGTATTAAGCTTTCCGCAGCTCAACGTGACCGACGGCGTCTACCACAGCGTGATAGTTCGCAGGCATGGA<br>GACTATGCGATCATGCAACTTGACTACAGCGGCTACGTCATCGGCAGCTTGCACAGTCAGCGAACCCTCCTGGACATGAGTGGTGGAGAGATAT<br>TTTCCGGGGGTCTCCCTAATATCACAATCGTCCGTATCATCGAAGCGATCGTTGAGAACGACGGGAGTGCTGTGATCAGCACGAACGTGCGTAA<br>CGATGGAGACGGTTACGCCGCCGACGTTGGAGGGGTGCATGTCCGAAATTTGCAATTGTCTGGGCCCCTTAACCTGGTGTCTCGTAAAAGGAG<br>AGCGTCTGGCACAGTCTCCGTGCTAGGAGACTTTGGAGGGTGTATAGCAGGGACCAGTGTGAATGGCGCCAACATGGAGAGTGACCCGTCCAT<br>CAAGGTCCATCGGCAGAACGTTCTAGACGGATGCCCATGTCTCTCTAATTTTTGCGCAAACGGAGGGACGTGCGTGGACGCTATGCCGCCCTAC<br>TGTATCTGTGCCCCAGGCTGGACCGGTCCACTGTGTACCATAGTGGTGACTGCCCCACCCGTCGGTGAGCGAGGCACGCCGTTCATGCACCCT<br>GCCGTCATCGCCATTATCTTAGTCGTCATGCTCGCCATCTTCATCATCATGGGCGCCGTCATCCTCAAGCGACGTCCTGAGCCCGTGGTCGTCTA<br>CGCTGACTCCACCGATACCGGCCACGTGCATGACAACGTGCGACTTTACCATGACGACGGCGGCGGGGAAGAGGACAATCTCGGCTACGACAT<br>CACCAAGCTGATGAAGTATACGTACATAGAGACCACTATAGCGCCGCCAAGTGTAGCGCCTTCTAAGGCATCCGAAGACAAGATATCTACCAGCT<br>CGGACCAGCCCCTGCTCCAAGGAAGGCCCCCGGATGCCGTGTTTGGCTTAACGGGCAAGGAACCAGGGCCCAAGATGCCCAAATACATGGAG<br>GGAGATGACGTGGGAGACTTTATCACAACACGAGTCAAAATAACCGATAGAGAGGTGTTCTTAGCTGTCGATGAACTGCACATCTACCGCTACGA<br>GGGAGACGACACAGACGTCGATGATCTGAGCGAGATCGAGCCAGACGAAGAGGATGAGGAATATGAACAAGAGTTCGATTTCCTCAAGCAGTG<br>GGGACCCAAGTTTGACAAACTTGCAAAGCTGTATGAAGACGTGGATGAGTAGTAATAGGGGATTTTTAAAATTGACATTTATGGAGGTAAAAAAGC<br>GGTTTTGAGAAGCACATCATTTCATGTCAGCAACATGCCACTCGATATCCTGATGGACTCGACGTGTGCAACGTTATAACGGCGACATGGAAGGT<br>GTCGTAGGCGTCACGTGGGGGATGCATGCAACTCAGCGAGCTATGTGACAATACGCGCATAGTGAACTGCTGGTTGCAGCGCAGTGTTGCTGA<br>ATTCTTCCTATTTACCAGAAGGCCATCCTAAAGAGTAATCCCATATAAAGTTAAAGTTCTATCATAACTAAGACCCTATGTATACATCCAAAAAATTCAT<br>TGTACATACAGACTTTCTTATCTAGAAACACCTATTTTTCCATGTATTAGTTCCTTTTTTATAGACGCTTTTAGGAATGATTTTGATGGCATTGTTTGAG<br>ATGTATAATGCTATGTGTTTTTTGCCTCTCGAACAGATTTTAAACAAACAAAAATATGCATGGCGTGTGCTTTTAAAACATTTACTGAATTTGCGTATT<br>AGATAGGGAATGTTAATAATTCGACTACTGTTGGTAAATGCGTGTGATGTCGGTTGAACTTTTAGCCACATTGACTTTGAAGCAAACGGCTATTAGT<br>CATGTGACAGTAATGATGTCACTGCACTGGTTTTTCATTTGCCCCCCCCCCCCCCCCCCCCCGATTGATTGACAGTTCATGCTTGTGCTTTTCA<br>GTCAAAGCAGTTTTCACCTTCAGACCATTTCACCCTATTCTCAATTAAATAAACTAAACCAACAAGTTAACGTAATTATGAATATTCTTGTGAATATTC<br>ATGAGTATTTCTTTCTACAGATTGATGCCCATAGTTTGGATTTCTCACTTTTAGATGGTTGCGGTGTGGCAATGCATCTTGGTCAGGGTGACTGGG<br>TAATCGATTAACTGTCCCTGTACTATTTAATTAGAGATGTCCATAATTTTGTTTAAAATAGAGAATATTTTCCAAGTAGCTTAGACTTGGTAAAGGATTG<br>GATGGACACGTATAATATTTTGTATTATTGTAGTTTTGTATTGTACAATGGTAAGATTTG |

**Table S2. HCR sequences**

| Group | singleR.labels | newIdents | avg_score | rank |
|---|---|---|---|---|
| Mesoderm_agg.4h | Mesoderm | agg.4h | 0,50482666 | 1 |
| Mesoderm_agg.12h | Mesoderm | agg.12h | 0,49179268 | 1 |
| Mesoderm_agg.48h | Mesoderm | agg.48h | 0,43979358 | 1 |
| Mesoderm_agg.24h | Mesoderm | agg.24h | 0,38139352 | 1 |
| Mesoderm_wt.24h | Mesoderm | wt.24h | 0,34019005 | 1 |
| Endoderm_agg.12h | Endoderm | agg.12h | 0,10119391 | 2 |
| Endoderm_agg.24h | Endoderm | agg.24h | 0,07551296 | 2 |
| Endoderm_agg.4h | Endoderm | agg.4h | 0,0665422 | 2 |
| Endoderm_agg.48h | Endoderm | agg.48h | 0,04158794 | 2 |
| Endoderm_wt.24h | Endoderm | wt.24h | -0,0092309 | 2 |
| Ectoderm_agg.12h | Ectoderm | agg.12h | 0,00679617 | 3 |
| Ectoderm_agg.4h | Ectoderm | agg.4h | 0,00313975 | 3 |
| Ectoderm_agg.24h | Ectoderm | agg.24h | -0,034524 | 3 |
| Ectoderm_agg.48h | Ectoderm | agg.48h | -0,0389231 | 3 |
| Ectoderm_wt.24h | Ectoderm | wt.24h | -0,0749227 | 3 |
| Neuronal_agg.4h | Neuronal | agg.4h | -0,0346418 | 4 |
| Gland Cells_agg.12h | Gland Cells | agg.12h | -0,0404037 | 4 |
| Neuronal_agg.24h | Neuronal | agg.24h | -0,0501314 | 4 |
| Neuronal_agg.48h | Neuronal | agg.48h | -0,0557535 | 4 |
| Secretory Progenitor_wt. | Secretory Progenitor | wt.24h | -0,0923819 | 4 |
| Secretory Progenitor_agg | Secretory Progenitor | agg.4h | -0,0385637 | 5 |
| Neuronal_agg.12h | Neuronal | agg.12h | -0,0515581 | 5 |
| Gland Cells_agg.48h | Gland Cells | agg.48h | -0,0564588 | 5 |
| Gland Cells_agg.24h | Gland Cells | agg.24h | -0,0645815 | 5 |
| Cnidocytes_wt.24h | Cnidocytes | wt.24h | -0,0982116 | 5 |
| Gland Cells_agg.4h | Gland Cells | agg.4h | -0,0498119 | 6 |
| Secretory Progenitor_agg | Secretory Progenitor | agg.12h | -0,0543732 | 6 |
| Secretory Progenitor_agg | Secretory Progenitor | agg.48h | -0,0755878 | 6 |
| Secretory Progenitor_agg | Secretory Progenitor | agg.24h | -0,0759984 | 6 |
| Neuronal_wt.24h | Neuronal | wt.24h | -0,0993192 | 6 |
| Cnidocytes_agg.12h | Cnidocytes | agg.12h | -0,0758632 | 7 |
| Cnidocytes_agg.24h | Cnidocytes | agg.24h | -0,0844906 | 7 |
| Cnidocytes_agg.4h | Cnidocytes | agg.4h | -0,0863492 | 7 |
| Cnidocytes_agg.48h | Cnidocytes | agg.48h | -0,0913767 | 7 |
| Gland Cells_wt.24h | Gland Cells | wt.24h | -0,103571 | 7 |

**Table S3. Cluster ranks based on projected module scores**

GAGGCGACGATGACTGTTCTATTGTGTTGGTCCTTTGATCTTAACTCTCCATGCTGGACGATCCCGAGCAGTGGTTTGCCAAG
AGTTTGAGCTTATATCCCTCCACTTCAGGTTCCTCTTTACCACATCTTTGAACCTGAGTCTAGGCCGACCCTGATTCCTGGTCC
CTGTCGTGAGCTGAGAGTAGAGCAGCTGCCTAGGCAATCTGTCAGTGTCCATGCGGTGAACATGCCCAGTCCATCTGAGATT
TTTCTCAATGAGTGTGTCAGCCATTGACCGTAATCCGGTCCTCTTGTATATCTCTTTGTTTGTGACTTTATCCATCCAAGTAATTT
TCATTATTTCTCTCAGGTGTCGCATCATATATGCATGTAAATTTTTCACCTGGCTGCGATATACAGTCCATGTTTCTGCACCGTAT
ACTAGGGCAGATAAGACAACTGCTCGATAGACCTTGCCTTTAACCTTGAGCGATACGTGATGGTTGTCCCACAGCCGCTCTCT
CAGCTTAGCATATGCTGCGCTGGCCTTCCCCTTTCTGAAGGTGATCTCATTATCTATTCGTGCATTGTCAGAGATGGTACTTCC
CAAGTACACAAAGTCTGGAAACAGTAGGGCGCTGAATACTCATTTTATCCTGAGAAACAGTCCATTCGACGTGAAAAGCTGAA
GTGATAAGAAATTTTGTTTCTTAATTTTCGAATAGTGTGAATTAAATGAAGTGAGGCCCGTATTTTTCAACTGGCTTGCTTGAATT
TTTAATTTCCATGCATACTTATTCTGACTAGTCTGTCAACATGGTTCTAAAACTTGAAAAAAATGATTTCCTTTTTTACAAGCAAAA
TCTGACCGACTTTCCCAAAAAAAGCCTCAAACGTGACGTTATATCGATCGGAGCTTATTAACCCATTGCCTCTTAGGCTGTACT
GCACCCTGTACCCTGTACTGCTCGTGGATAAATTAAACAAGAGCCTCCAAAAATCATAAATGCAAATATAACAACTAGATCCAGA
TACTCAGGCATTTAGCCTATGGGATATTAGGCTTTTTTTTAAATATACAACCCAACTGTGAGATGAACTACTACTGTACAGCCAAA
TATTGTAGCTCAAGTTCTTCGACAAATTCCCAACTAGACAAGAAAAACATGTTCGAGTCACAAGTTTTTATCTTTATTTTGCCCA
CCAGGTTCAGCAATCTCATTGAAACTTATTTGTAAAATTTAAACTTTAAAAACTTAATAAAGTTTTTATCTTTATTTTGCCCACCAG
GTTCAGCAATCTCATTGAAACTTATTTGTAAAATTTAAACTTTAAAAACTTAGTATTCTTTAAAGCTCATCTGTGGATCGCGCTAAG
TATACGCGCGTCTCGTCGCCGTCGAAATGAGGCCAGATAGAAAAATCCACAAGTCAAAAAAGGTCAAGTCAAAAAGCCGCGC
AATGTTTTCCAGAGCTTGCATTCTTCAATTTCAGATTGCGCTTTGTGCATTGTTTTTTCGATAATGCTGTGCTTAATCAGACACC
CACAAAGTAGTCAAAGTGTTTTTGTTTTAAACTGCAACAGTATTTAATTATTATTGGGTGTGTATTCGTAGATGAGACATTGATGT
TGAGAGCTATCTGTATAACCGCTTTTTTTCCCGTTGCAGCTCATTGTTTGTATTGTTTGCAAGGTGATAGGGCGTGTTTAGGTAA
ACCGGGGTCAACATCCGGGAGTCCGTTAAACACGGAAGCCCTCTTCAGCAACCCGCCAATCAACTGATCTTATTTATACTAGC
CTGACTTATCTGGCTAATAAGTCTTAACCAATGTATTGATAACGTCTTAAATCCAGCACTGAAAGCGCCAAACATATTTATAGAGC
TTGAAAGAAACATATAAGAAATACTCGTGACCGCGAGAGGGTCTCGATTCACGCAAACATGAATGCCGTCCCGTTCGACCAGT
GGAGAAACACATCCTGACAGTTCACGGGAAACGATGACACAAATCTTGCGTGACATCACTCCATTTGAATTAGATAGATATATC
GTAGACGATCTGTAGAGTTTACAAGTCGATACTTTCCGCACAGTTTAAGTCAAGAGCAAAGTAAATTCTAAGAGTTATTACCTTT
AGTTGTATCGTCAAACCAATTTGTCGCCAGTAAGAGGATAATGTTGTGTGGTAAGTGATCCGCTATTTATTGATCAAGTTTCTTA
AAAACTCGCTAAACTTGCCTCAGGGTGAAGTCAGTGAGATTACCTACCAACAGAACTCAGATGATTACGGAAAGTCCATACCT
CTTGGGTGTGTCAAAATAAAATGCGCCACTAAGAGATTCTTACTCTGTGCTTTGTGGGGAAAGACACGCAAGATACCCTCCTC
GCTTCTTATTTGATTAGTATGTAACTCTAGACCCGAGTTGTAAATGTAAGCTAACACCCTACCCTCCCAAGGACCAAGATCTTCG
TGGGGAAATCCTCTAGTAGATTTGACGCCATCCGATTGGCTGGTAGTCGCATTGCTAACATTCATTGACGCGTATCGTGTTTCC
GGTTCACTCTTGCTGCCTTCCACTCCTTGTAACTTGGGCTTTACGCCAGTCCGTTTGGTCCTAAGTGTGAGGATTCTGCAACA
ATACCGGTGCAGTGGAGTCGAGGGAGGTAGCTAGGGCGTCCTGGAAGTGCTACTGGACTTGCTGGCGTAGAATTAGCTTGTT
GGACTGGTTGTTTTGTGAGGGAACCCCGTCATATAGCAAGCCTTGTCTGGTCGATCGCTTAGGAAGTCTGACTTCAATATTGC
AGCAAATCCTACATTATATTTTACCACTGTGTCCGTGAGTGAATTTTGAAAGGTTTGTTTTTCTGCCTACAACGACTCGACTGCT
TATTTTGGGCTAAGGTTTGTCTATCTCGAAAACAAAGTGGTCTTTGTGGTGGTAAATGCTTATATACTGTGCCGGCTATGGGTAA
CAGACCCTAAGCTGAAATATCAATTCATCAGATAACTCTCTTCAATGAATTGCTTTCTCAGTTCGGTTTTGCATCCGTCACACAA
GTCTTGTAACTTATAGTTTGCCTGATATTATTACAGATAAACAGATAAAGCTCAACAGTGGAAATACCCAAAGGGAAAATTTGAGT
TGTTTACGAATTAGAAACCCACACAAGACGATTCAGCCCGGCCATCCTCAGGTGTTTCACAACCTTCACCATGTTATTCGGTAA
GTAAACCTGCTACGCTTTCTATGTTATTTTGTTGGTTATATGACTCCTTAATTTTTTCAACCTGTCCTCTGAGTGGAAATATTTTTT
TCGTGCTTTGTTTACAAAATTCCAGGTATCACCGTTTATCGATCTTGAAACGCGAGTAAAAAATTCCTTGGATATTTTCTATTCCG
TGGTTTTCCATCCCTAAATCAAACGTTCATCAATGACTCGAGAATACATAATTCTAACCAGACAGTAGACAACTTAAGATAGATTA
CTAAACCCGATAATTATATAGTTGAACTTTTATCTGCGCTTGCTCTCAAATGACGTTTCGTCTTCTTGCTATTACTGTCACTGTTTT
CCTCCCGTGACCGCTCTAGTACTTCAACAAACACCGATATCTTATATCTTACGTTTGTAAAGCATGGGAAATCACGATTCTATGG
ATAAGGCCATAAGGAATGCCATATAATTCCTATAATAGCGCTGAAGTATGGTGAATATCACGTTATGCAATCTTGACCACATAAGG
CGGCATATCATGTCTCCATCTCGTTATTTTACACAAGTGAGTCTCGCTATATTTTAGAGACCTCCCGGCCTGATTCTGGCTCAAG
TATATGTTTGTTTGTTTTTACGAAAAATGAAAAAAATGTTAGGGGAGCCGGAACTAAGTTGAAACTATTCACTAACCAATCAAATG
GCACTATGTTTCGGGTTCCGGAATAAGCGGCTCCGGAATATAAAGTTGTAAAACAACTTGTGACTGATATTGAATTTCCCCATAA
CTTCTCATTAATGATTTGAGCAAGCTGACGCTCTATTCTCTGAATTGTGTTACTCAAGCATAAAATATACTCAACAGTAACCCAGT
ACGCGCTGACGAATATGTCCAGGAGGATAAACAATTCGATATAATTTTTTTACATGAGTACTTTAGATTTAAACCGTCAAGCCTTG
ACAAGGTCTGATGTCATGCACTTAAACCCCAGAGAACCTGCAAAGATGTTACCCCAGATGATTGTTTTAGTACTCATCTAAGTG
CACAGGTCGATAACTGTGTATGAAGCCCTAGGGAGGGAAAACACAGAAGTGCGCGCGTACATATGGGACCGCAAAGAAATCA
TGGAGGGTTGAGGGATTAGGTGACAGCCAAAAACCCTTCCCAAAAATCTTCATGTTTATCAGGCCCTAGGGAGGGAAAACAC
AGAAGTGCGCGCGTACATATGGGACCGCAAAGAAATCATGGAGGGTTGAGGGATTAGGTGACAGCCAAAAACCCTTCCCAAA
AAATCTTCATGTTTATCAGGCCCTAGGGAGGGAAAACACAGAAGTGCGCGCGTACATATGGGACCGCAAAGAAATCATGGAG
GGTTGAGGGATTAGGTGACAGCCCTACCCCGCAATGTCCAATGCTTCATGTTTCCGATTTTCAAGCTAGCTCACTAACACTTCC
AATGATCCATGTCTCCTTAG

**Table S4. Cloned promoter sequence of Betalaminin**
